# Different ways of evolving tool-using brains in teleosts and amniotes

**DOI:** 10.1101/2022.11.04.515163

**Authors:** Pierre Estienne, Matthieu Simion, Hanako Hagio, Naoyuki Yamamoto, Arnim Jenett, Kei Yamamoto

## Abstract

In mammals and birds, tool-using species are characterized by a high degree of encephalization with a relatively large telencephalon containing a higher proportion of total brain neurons compared to other species. Some teleost species in the wrasse family have convergently evolved tool-using abilities. In this study, we compared the brains of tool-using wrasses with various teleost species from a broad phylogenetic range. Using the isotropic fractionator, we show that in the tool-using wrasses, the telencephalon and the ventral part of the forebrain and midbrain are significantly enlarged compared to other teleost species but do not contain a larger proportion of cells. Instead, we found with tract tracing and selective neuronal fiber visualization that this size difference is due to large fiber tracts connecting the dorsal part of the telencephalon (pallium) to the inferior lobe (IL), a ventral mesencephalic structure absent in amniotes. The high degree of connectivity between the IL and the pallium in tool-using wrasses suggests that this unique teleostean structure could contribute to higher-order cognitive functions. Given remarkable differences in their overall brain organization, we conclude that, unlike in amniotes, the evolution of non-telencephalic structures might have been key in the emergence of higher-order cognitive functions in teleosts.

## Introduction

In primates, the cerebral cortex is the center for higher-order cognition such as logical thinking or self-recognition. However, some birds, such as corvids and parrots, demonstrate comparable cognitive functions, even though they do not possess this six-layered cortical structure (Emery and Clayton, 2004; Kirsch et al., 2008). Remarkable behaviors indicative of so-called higher-order cognition include tool use (Auersperg et al., 2012; Hunt, 1996; Hunt and Gray, 2004) and mirror use (Prior et al., 2008), which require causal reasoning, planning, or self-recognition.

Encephalization, the increased relative mass of the brain compared to body mass (Tsuboi et al., 2018), has long been used as a proxy for intelligence in vertebrates, with highly encephalized species considered more intelligent (Benson-Amram et al., 2016; MacLean et al., 2014). Despite the high degree of encephalization in corvids and parrots, cognitive abilities in birds have long been underestimated due to their rather small brains compared to mammals. A more recent cell counting study has in fact revealed that the brains of parrots and songbirds are extremely neuron-dense and contain on average twice as many neurons as primate brains of the same mass (Olkowicz et al., 2016). Thus, some species of corvids and parrots have as many neurons in their pallium (the dorsal telencephalon that contains the cerebral cortex in mammals) as some species of primates (Olkowicz et al., 2016). This strongly suggests that mammals and birds have followed two independent trajectories of encephalization: an increase in cortical surface in mammals (with the cortex reaching a very large size in humans), and an increase in the neuronal density of the pallium in birds. Both trajectories led to an increase in the absolute number of telencephalic neurons in highly encephalized species of mammals and birds compared to poorly encephalized ones. In other words, encephalization in amniotes (the clade containing mammals and birds) is mostly a process of “telencephalization” (Olkowicz et al., 2016; Ströckens et al., 2022; von Eugen et al., 2020).

Teleost brains are generally much less encephalized compared to amniotes (Tsuboi et al., 2018). Nonetheless, some teleost fishes belonging to the family of wrasses (*Labridae*) exhibit tool use-like behavior (Brown, 2012) or mirror use behavior (Kohda et al., 2019), that are observed only in a few species of mammals and birds. Thus, the brain anatomy of tool-using fish is of great interest to understand the evolution of higher-order cognitive functions. Our developmental studies have shown that the telencephalon, hypothalamus, and sensory nuclei of teleosts (Bloch et al., 2020; Fontaine et al., 2015; Xavier et al., 2017; K. Yamamoto et al., 2017; K. Yamamoto and Bloch, 2017) differ greatly from amniotes. Indeed, teleost brain organization appears to be much less conserved than previously thought. Notably, teleosts possess a remarkable ventral structure called the inferior lobe (IL), which is absent in tetrapod brains and whose functions remain largely unknown (Bloch et al., 2019). At a gross morphological level, the ventral parts of the teleost brain appear much more developed in comparison to amniote brains.

These observations raise the question of how encephalization occurred in the teleost lineage, and how the brains of teleosts with remarkable cognitive abilities, such as wrasses, may differ from other species. Is the teleost pallium the brain center that is responsible for higher-order cognitive functions, similar to the amniote brain? In order to uncover which brain structures are expanded in teleost species demonstrating complex behavioral repertoires, we examined the cellular composition of their major brain regions and compared them with other teleost species located at various phylogenetic positions.

Our quantitative study revealed that, unlike in amniotes, the ventral part of the brain including the IL is significantly developed in the brains of tool-using wrasses, illustrating how encephalization in teleosts and amniotes followed different evolutionary paths.

## Results

### The tool-using wrasse *Choerodon anchorago* has more cells in its brain than the hamster

The body mass and brain mass were measured for 11 species of teleost: a group of 3 wrasse species (*Choerodon anchorago*, *Labroides dimidiatus, Thalassoma hardwicke*), a group of 4 cichlid species (*Maylandia zebra*, *Neolamprologus brichardi*, *Ophthalmotilapia boops*, *Amatitlania nigrofasciata*), and a group of 4 other species (the medaka (*Oryzias latipes*), zebrafish (*Danio rerio*), *Astyanax* surface fish (*Astyanax mexicanus*), and trout (*Salmo trutta*), hereafter referred to as the “outgroup”) (Figure 1). Total number of cells in the brain was determined using the isotropic fractionator (see “Materials and Methods” section).

**Figure 1.**
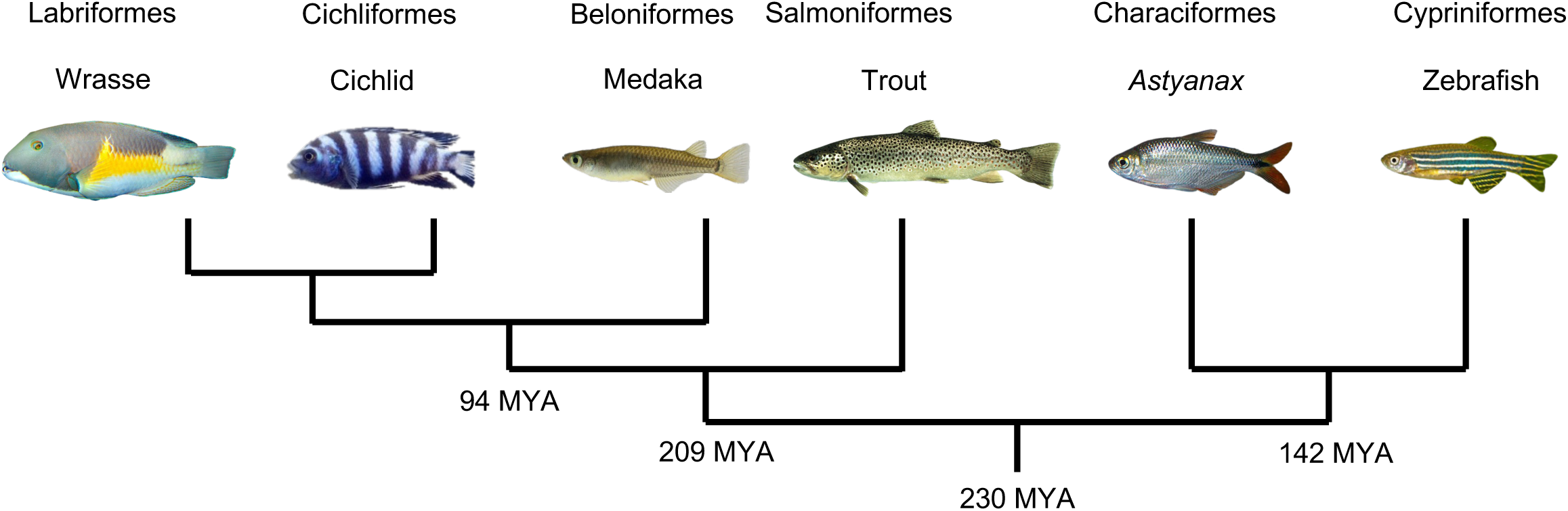
Phylogenetic tree of the teleost species sampled in this study. The families of wrasses and cichlids are the closest relatives within teleosts. In this study, we refer to the medaka, trout, *Astyanax*, and zebrafish as the “outgroup”. Numbers at the root of each tree branches represent the estimated time of divergence (MYA: million years ago). The last common ancestor of these species can be traced back to 230 million years ago (http://www.timetree.org (Kumar et al., 2022)).

Remarkably complex behaviors (tool use in the case of *C. anchorago* and *T. hardwicke* (Bernardi, 2012; Paśko, 2010), and social cognition in the case of *L. dimidiatus* (Kohda et al., 2022; McAuliffe et al., 2021; Wismer et al., 2016)) have been reported in the group of wrasses. Cichlids are phylogenetically close to wrasses and display relatively complex social and cognitive behaviors, although no instances of tool use have been observed (Barks and Godin, 2013; Hotta et al., 2019; Schluessel et al., 2022; Snekser and Itzkowitz, 2020). By contrast, no such behaviors have been reported in the “outgroup” species.

Among the wrasses studied, body mass ranged from 1.55 to 91.52 g, brain mass from 34.62 to 338.8 mg, and total number of cells in the brain from 45.7 to 185.08 million (Table 1). Wrasses were wild caught and were generally young adults; however, one large adult of *C. anchorago*, weighing around ten times as much as the other individuals, was also sampled. Statistical analysis showed that the data from this large individual did not impact the statistical significance of our results (see Supplementary file 1, Figure 2-figure supplement 1, Figure 3-figure supplements 1 & 3). In cichlids, body mass ranged from 5.15 to 20.28 g, brain mass from 42.43 to 96.94 mg and total number of cells in the brain from 37.54 to 61.78 million (Table 1). In the “outgroup”, body mass ranged from 0.492 to 177.15 g, brain mass from 8.38 to 354.73 mg, and total number of cells in the brain from 6.66 to 100.84 million (Table 1).

**Figure 2.**
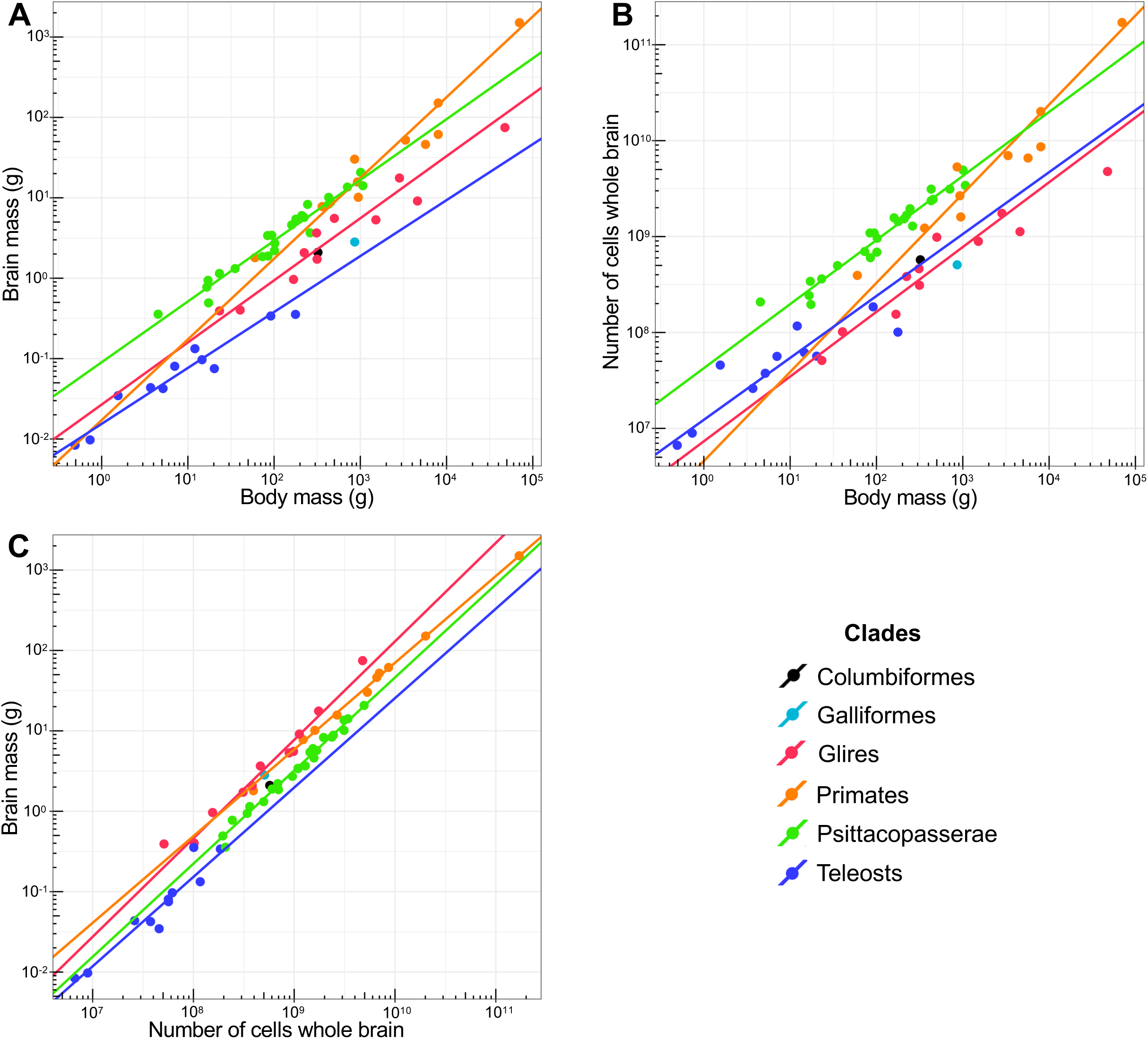
Teleosts have small, cell-dense brains that contain more cells than the brains of rodents of similar body mass. The fitted reduced major axis (RMA) regression lines are displayed only for correlations that are significant (P<0.05). Each point represents the mean value of a species. X and y axes are in log^10^ scales. Body mass and Brain mass, Total number of cells in the brain and Brain mass, and Body mass and Total number of cells in the brain were significantly correlated in all groups (Spearman r ranging from 0.945 to 1; p<0.0001 in all cases). Data on Columbiformes and Galliformes (Olkowicz et al., 2016) was plotted as illustration but wasn’t included in the statistical analysis due to the small sample size. All regression lines are significantly different, except for the regression lines of Glires and Primates in plot (C). (A) Brain mass plotted as a function of body mass. Teleosts have smaller brains than birds and mammals of similar body mass. Regression lines for Body mass and Brain mass had significantly different slopes (ANCOVA, p<0.0001). Pairwise comparisons found significant differences in the slopes of Glires and Primates (p<0.001), Primates and Psittacopasserae (p=0.0001) and Primates and Teleosts (p<0.0001). ANCOVA revealed significant differences in the intercepts of the regression lines for Brain mass and Body mass for groups with statistically homogenous slopes (p<0.0001). Pairwise comparisons found significant differences in the intercepts of Glires, Teleosts and Psittacopasserae (p<0.0001 in all cases). (B) Total number of cells in the brain plotted as a function of body mass. Teleost brains contain less cells than bird and primate brains, but more cells than the brains of rodents of similar body mass. Regression lines for Body mass and Total number of cells in the brain had significantly different slopes (ANCOVA, p<0.0001). Pairwise comparisons found significant differences in the slopes of Glires and Primates (p<0.001), Primates and Psittacopasserae (p<0.0001) and Primates and Teleosts (p<0.001). ANCOVA revealed significant differences in the intercepts of the regression lines for Body mass and Total number of cells in the brain for groups with statistically homogenous slopes (p<0.0001). Pairwise comparisons found significant differences in the intercepts of Glires, Teleosts and Psittacopasserae (p<0.05 in all cases). (C) Brain mass plotted as a function of total number of cells in the brain. Cellular density inside the teleost brain is higher than in birds and mammals. Regression lines for Total number of cells in the brain and Brain mass had significantly different slopes (ANCOVA, p<0.0001). Pairwise comparisons found significant differences in the slopes of Glires and Primates (p<0.01) and Primates and Psittacopasserae (p<0.01). ANCOVA revealed significant differences in the intercepts of the regression lines (p<0.0001). Pairwise comparisons found significant differences in the intercepts of the regression lines for Glires, Teleosts, Primates and Psittacopasserae (p<0.0001 in all cases), with the exception of the intercepts of Glires and Primates (p=0.08). Glires = rodents and lagomorphs. Psittacopasserae = Passeriformes (songbirds) and Psittaciformes (parrots). See also Table 1.

**Figure 3.**
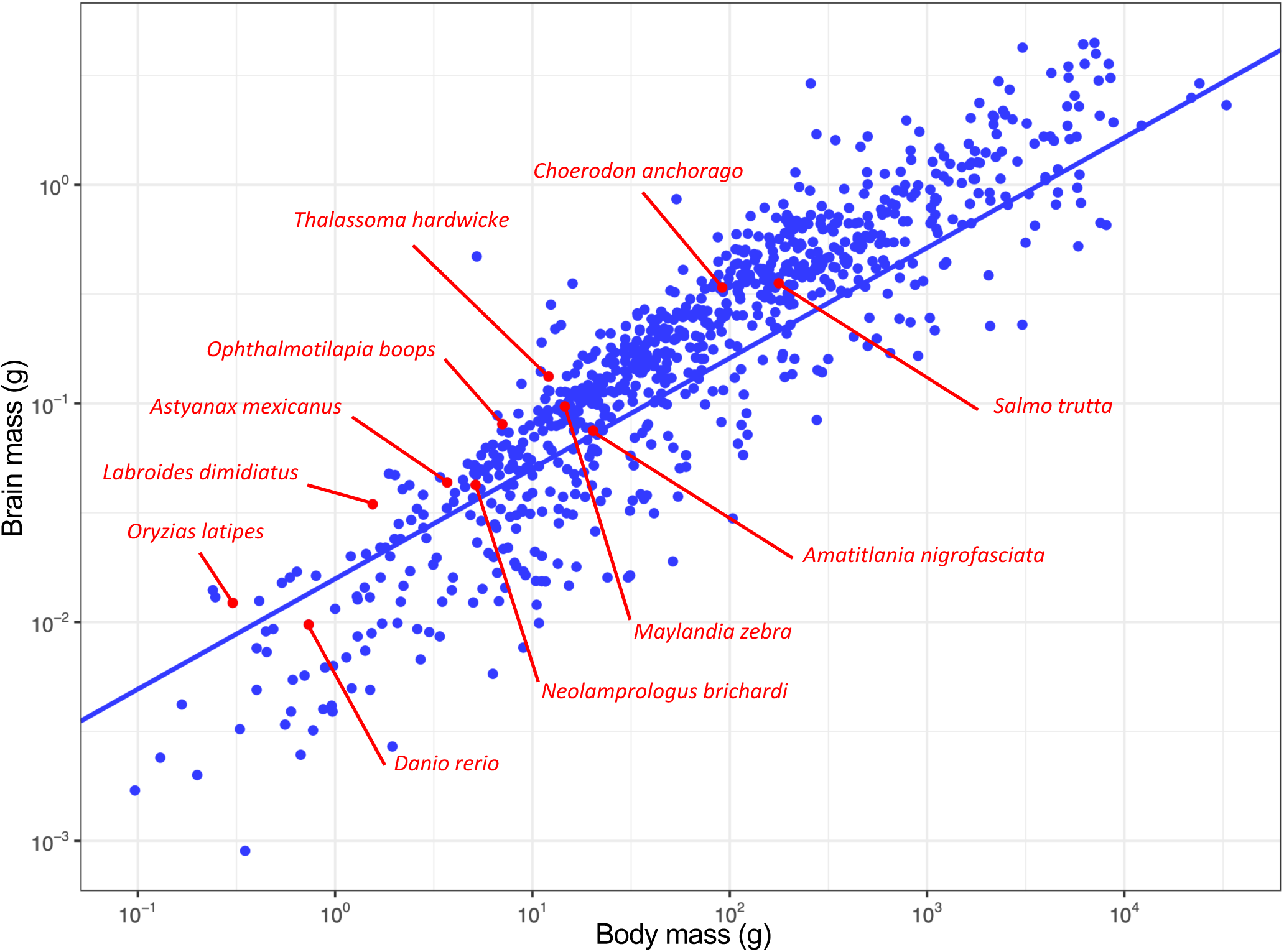
Encephalization in 11 species of teleosts (red) compared to a large dataset of actinopterygians (blue). Brain mass is plotted as a function of body mass, and the phylogenetically corrected (phylogenetically generalized least squares regression test, PGLS) allometric line is shown. Each point represents the mean value of a species. X and y axes are in log^10^ scales. The phylogenetic regression slope for actinopterygians is of 0.50 ± 0.01. Adjusted R^2^: 0.8382, t=65.978, p<0.0001. See also Table 2.

**Table 1.** Cellular composition of the brains of 11 teleost species. All values are mean ± SD.

| Species | n | Body mass (g) | Brain mass (mg) | Total cells ($\times 10^6$) |
| --- | --- | --- | --- | --- |
| **Wrasses** |  |  |  |  |
| *Labroides dimidiatus* | 3 | 1.55 ± 0.35 | 34.62 ± 5.62 | 45.7 ± 6.67 |
| *Thalassoma hardwicke* | 3 | 12.07 ± 4.14 | 132.82 ± 23.49 | 116.78 ± 25.62 |
| *Choerodon anchorago* | 4 | 91.52 ± 137.39 | 338.80 ± 206.95 | 185.08 ± 78.73 |
| *Choerodon anchorago* * | 3 | 22.85 ± 4.54 | 235.44 ± 11.67 | 146.11 ± 13.59 |
| **Cichlids** |  |  |  |  |
| *Neolamprologus brichardi* | 5 | 5.15 ± 1.53 | 42.43 ± 4.26 | 37.54 ± 7.08 |
| *Amatitlania nigrofasciata* | 5 | 20.28 ± 7.85 | 75.05 ± 12.4 | 56.62 ± 6.37 |
| *Ophthalmotilapia boops* | 3 | 7.04 ± 2.39 | 80.32 ± 7.27 | 56.37 ± 6.35 |
| *Maylandia zebra* | 3 | 14.59 ± 2.58 | 96.94 ± 6.62 | 61.78 ± 3.72 |
| **Others** |  |  |  |  |
| *Oryzias latipes* | 5 | 0.492 ± 0.07 | 8.38 ± 1.12 | 6.66 ± 0.54 |
| *Danio rerio* | 5 | 0.73 ± 0.21 | 9.74 ± 0.2 | 8.92 ± 0.69 |
| *Astyanax mexicanus* | 5 | 3.7 ± 1.17 | 43.55 ± 4.35 | 26.09 ± 2.74 |
| *Salmo trutta* | 4 | 177.15 ± 45.05 | 354.73 ± 33.16 | 100.84 ± 8.58 |

*Choerodon anchorago*\* = large individual of *Choerodon anchorago* excluded.

**Table 2.** Residuals obtained by fitting a log^10^-log^10^ regression of brain mass against body mass for 11 species of teleost. All values are rounded to the nearest ten thousandths.

| **Species** | **Residual** | **Residual excluding the large *C. anchorago* individual** |
| --- | --- | --- |
| *Thalassoma hardwicke* | 0.3797 | 0.3792 |
| *Choerodon anchorago* | 0.3420 | 0.4879 |
| *Ophthalmotilapia boops* | 0.2795 | 0.2790 |
| *Labroides dimidiatus* | 0.2458 | 0.2451 |
| *Salmo trutta* | 0.2171 | 0.2170 |
| *Maylandia zebra* | 0.2012 | 0.2008 |
| *Astyanax mexicanus* | 0.1549 | 0.1543 |
| *Oryzias latipes* | 0.1530 | 0.1520 |
| *Neolamprologus brichardi* | 0.0708 | 0.0703 |
| *Amatitlania nigrofasciata* | 0.0179 | 0.0176 |
| *Danio rerio* | -0.1409 | -0.1417 |

Compared to previously published data, teleosts had smaller brains than birds, primates or rodents of similar body mass (Figure 2A). By contrast, teleost brains contained more cells than the brains of rodents of similar body mass, albeit not as many as birds and primates (Figure 2B). For instance, the brain of the tool-using wrasse *C. anchorago* contained on average more cells than the brain of the nearly 2 times heavier hamster (*Cricetus cricetus*).

Cellular density inside the teleost brain was higher than in birds and mammals, with teleosts having as many cells as rodent brains more than 4 times larger (Figure 2C). For example, the large *C. anchorago* individual sampled had 301.9 million cells in its brain, nearly as many cells as a rat (*Rattus norvegicus*), even though its brain was 2.6 times smaller.

### Encephalization and relative mass or number of cells in the telencephalon of teleosts are not correlated

Residuals obtained by fitting a log^10^-log^10^ regression of brain mass against body mass data from this dataset with previously published data on actinopterygians (Figure 3) using a phylogenetic generalized least square (PGLS) model ranged from -0.142 (*D. rerio*) to 0.38 (the wrasse *T. hardwicke*), with only one other species (*C. anchorago*) with a residual >0.30 (Figure 3, Table 2). Excluding the large *C. anchorago* from our analysis gave a residual of 0.488 for *C. anchorago*, placing it above *T. hardwicke* (Table 2, see Supplementary file 1). Overall, these two tool-using species were the most encephalized of our dataset.

In order to compare the degree of encephalization with the relative mass and cellular composition of major brain regions, the brains of ten species were dissected into five parts (Figure 4A): the telencephalon (Tel), the optic tectum (TeO), the rest of the Forebrain/Midbrain (rFM), the cerebellum (Cb) and the rest of the Hindbrain (rH) following the rostro-caudal and dorso-ventral axis (Figure 4B-E, See “Materials and Methods” section). These structures were weighed and the number of cells contained in each structure was determined using the isotropic fractionator. No statistically significant correlations were found between encephalization and the relative mass and relative number of cells of the Tel, TeO, rFM, Cb (Figure 3-figure supplement 2A-D). The only structure that showed a statistically significant correlation with encephalization was the rH (Figure 3-figure supplement 2E), with a negative correlation for both the relative mass and relative number of cells. This indicates that more encephalized species of teleosts have a relatively smaller rH containing a relatively smaller number of cells.

**Figure 4.**
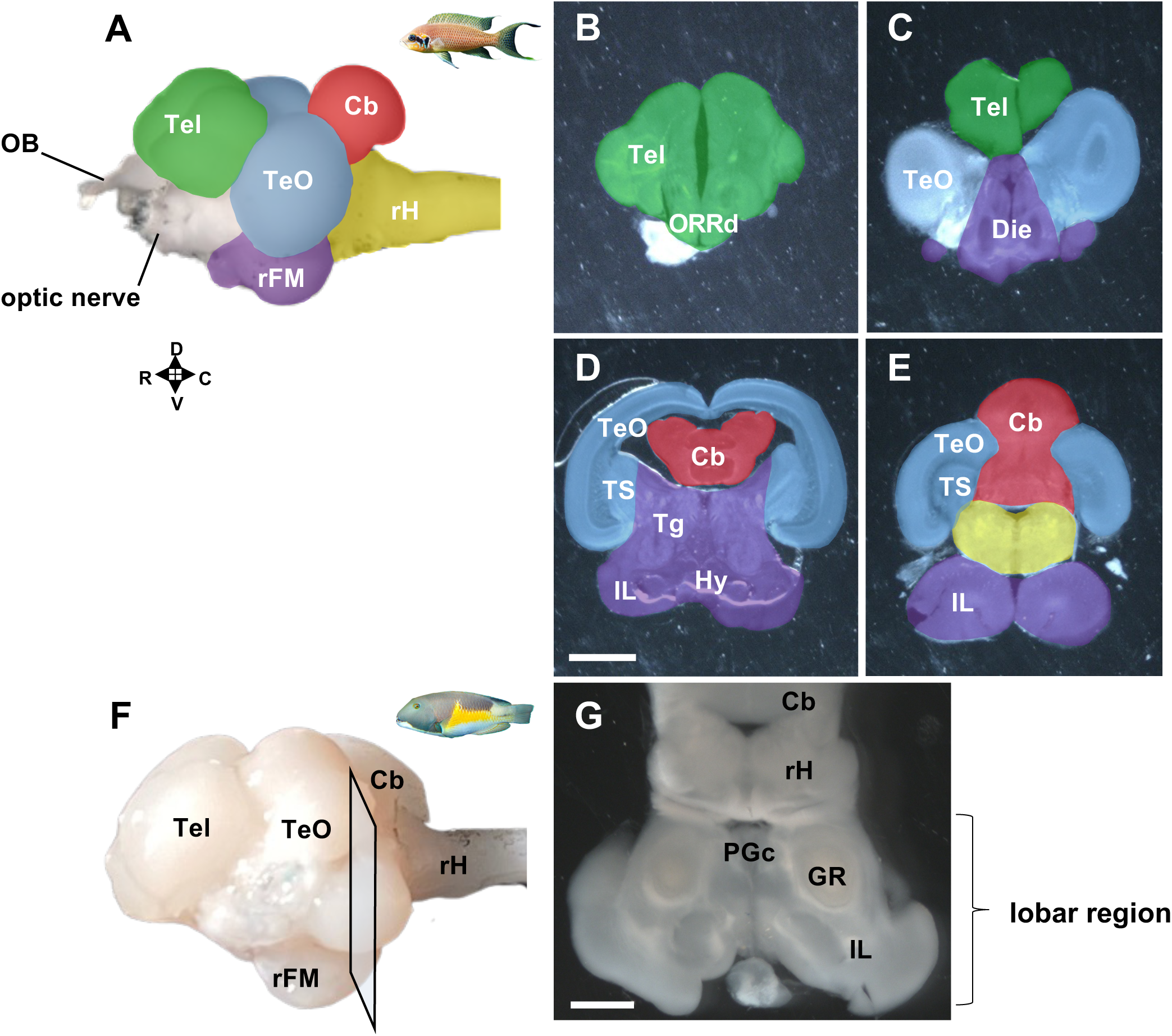
Illustration of brain structures of teleosts. (A-E) Dissection of the five major structures for the isotropic fractionator. (A) Lateral external view of the brain of the cichlid *Neolamprologus brichardi*. The different brain regions are color-coded. The uncolored regions are the olfactory bulbs and cranial nerves. (B-E) 300 µm frontal sections of the brain of *Neolamprologus brichardi* from rostral to caudal, showing the boundaries of the five major brain regions. The regions are highlighted following the color code in (A). (F,G) Illustration of the lobar region. (F) Lateral external view of the brain of the wrasse *Choerodon anchorago* indicating the level of the frontal section shown in (G). We collectively refer to the area containing the PGc, GR, and IL as the lobar region, which is a teleost-specific structure absent in the tetrapod brain. Brain regions: Cb: cerebellum; Die: diencephalon; GR: corpus glomerulosum pars rotunda; Hy: hypothalamus; IL: inferior lobe; ORRd: dorsal optic recess region; PGc; preglomerular nucleus pars commisuralis; rFM: rest of the forebrain/midbrain; rH: rest of the hindrain; Tel: telencephalon; Tg: tegmentum; TeO: optic tectum; TS: torus semicircularis. Scale bars: 1 mm. R: rostral; C: caudal; D: dorsal; V: ventral. See also “Materials and Methods” section.

These results suggest that teleost brains have evolved very differently from amniote brains. Unlike in mammals and birds, encephalized species of teleosts don’t have an extremely large Tel.

### Wrasses have a relatively larger Tel and rFM, but not a larger relative number of cells compared to other teleosts

Comparing species in a one-to-one manner didn’t reveal any consistent differences in either the relative mass or relative number of cells across structures (Figure 5-figure supplement 1). However, a trend towards larger Tel and rFM was observed when examining wrasses as a whole (Figure 5-figure supplement 1). Wrasses have large brains and display the most flexible behavioral repertoires, including tool use. We thus aimed to investigate any differences in their brain morphology compared to the other teleosts. To this end, the relative mass and number of cells in the five major regions of the brains of all wrasse species were compared with those of the other species of teleosts sampled in this study (Figure 5).

**Figure 5.**
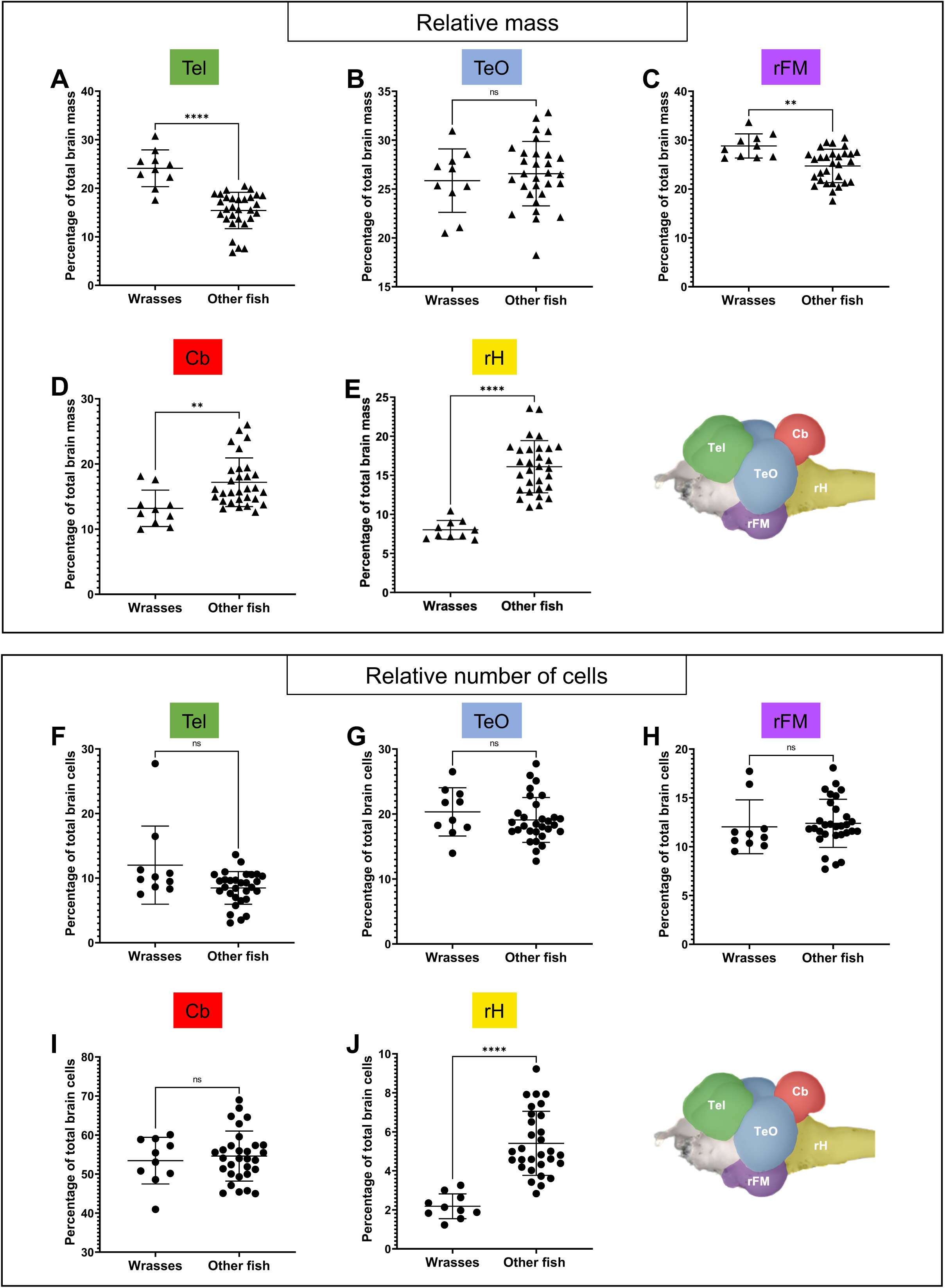
Relatively larger Tel and rFM without a corresponding increase in the relative number of cells in wrasses compared to other teleosts. Three species of wrasses (« Wrasses »: *Choerodon anchorago*, *Labroides dimidiatus*, *Thalassoma hardwicke*, n=10) were compared with seven species of teleosts of various orders (« Other fish »: *Astyanax* mexicanus, *Amatitlania nigrofasciata, Danio rerio, Maylandia zebra, Neolamprologus brichardi, Ophtalmotilapia boops, Salmo trutta*, n=30). Top panel (A-E): Comparison of the relative mass of the Tel (A), TeO (B), rFM (C), Cb (D), and rH (E). Wrasses have a relatively larger Tel and rFM compared to other teleosts. Bottom panel (F-J): Comparison of the relative number of cells in the Tel (F), TeO (G), rFM (H), Cb (I), and rH (J). Despite having a relatively larger Tel and rFM, wrasses don’t have a larger proportion of cells in those structures compared to other teleosts. Statistical analysis: Mann-Whitney’s test. Each point represents individual values. Error bars: mean ± SD. ns: non significant, **p<0.01, ****p<0.0001. Brain regions: Cb: cerebellum, rFM: rest of the forebrain/midbrain; rH: rest of the hindrain; Tel: telencephalon; TeO: optic tectum.

The relative mass of the Tel and rFM was significantly higher in wrasses compared to other teleosts, with the Tel accounting for 24.11% ± 3.78% of total brain mass in wrasses compared to 15.43% ± 3.74% in other species (Figure 5A). While the Tel in wrasses was larger than in other teleosts, it remained modest when compared to amniotes. The rFM accounted for 28.82% ± 2.46% of total brain mass in wrasses compared to 24.72% ± 3.43% in other species (Figure 5C). The relative mass of the Cb and rH were significantly lower in wrasses compared to other teleosts (Figure 5D-E). No significant difference was found in the relative mass of the TeO between the two groups (Figure 5B).

Despite the larger size of the Tel and rFM in wrasses, isotropic fractionator data revealed that there was no significant difference in the relative number of cells in these two structures compared to other species. The Tel accounted for 12.02% ± 6.04% of total brain cells in wrasses compared to 8.49% ± 2.54% in other species (Figure 5F), while the rFM accounted for 12.04% ± 2.76% of total brain cells in wrasses and 12.39% ± 2.47% in other species (Figure 5H). No significant difference in relative number of cells was found in either the Cb (Figure 5I) or TeO (Figure 5G), whereas the rH (Figure 5J) accounted for a significantly smaller relative number of cells in wrasses compared to other species (2.18% ± 0.64% and 5.41% ± 1.64%, respectively).

Similar results were obtained when comparing the group formed by wrasses and cichlids together to the “outgroup” (Figure 5-figure supplements 2 & 3).

Overall, these results show that wrasses have a relatively larger Tel and rFM compared to other teleosts. However, these two structures do not contain a larger proportion of cells than in other teleosts.

### Pallium and IL display increased connectivity in wrasses compared to other teleosts

We hypothesized that the increase in mass observed in the Tel and rFM of wrasses was due to an increase in the neuropil of these structures. To verify this hypothesis, we performed selective visualization of the fibers in the Tel and rFM. Whole brains of the wrasse *C. anchorago*, the cichlid *N. brichardi*, the trout *S. trutta*, the *Astyanax* surface fish *A. mexicanus*, and the zebrafish *D. rerio* were cleared, stained with DiI, and imaged on a light-sheet microscope (Figure 6, Videos 1-5, See «Materials and Methods» section).

**Figure 6.**
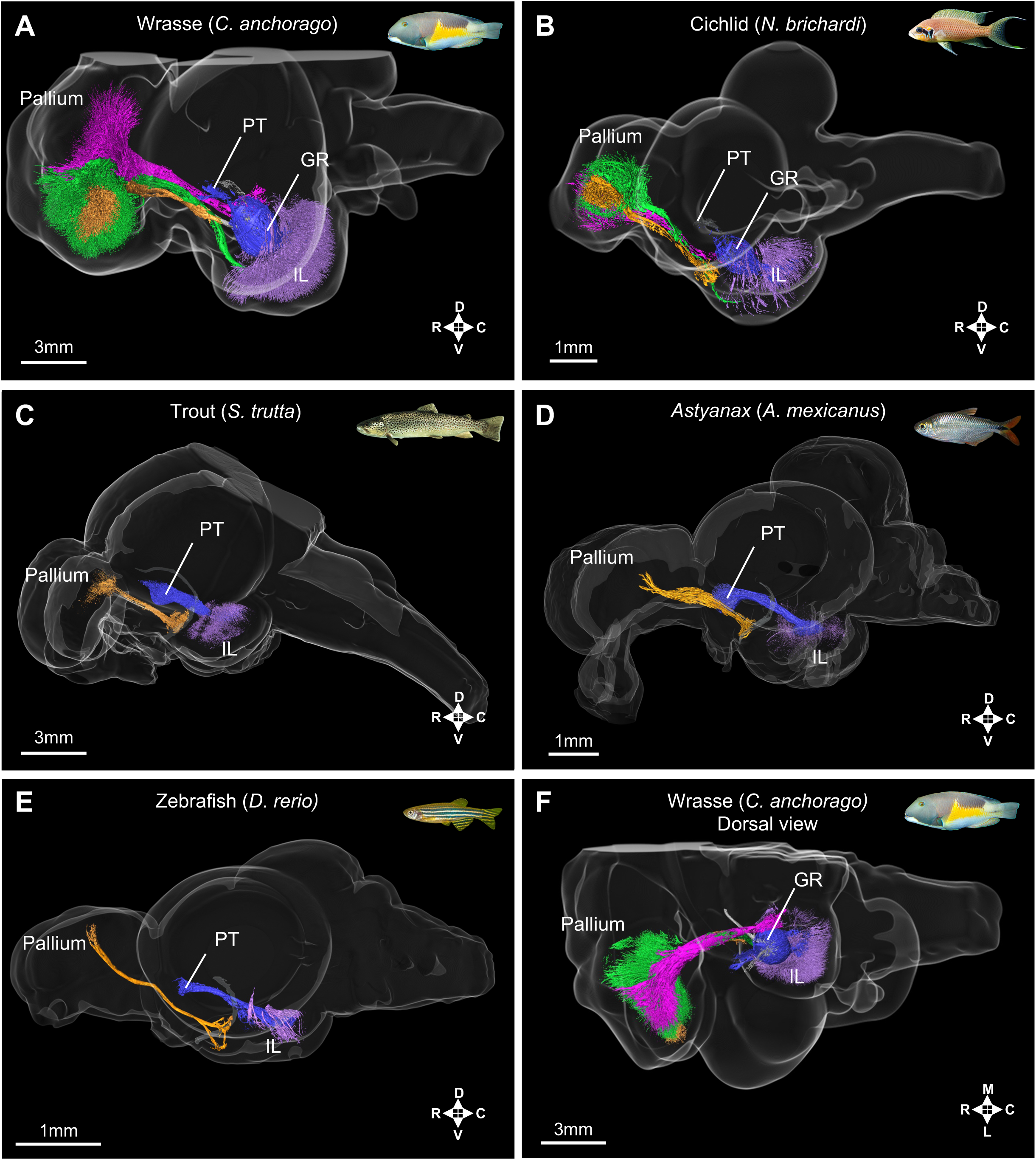
The pallio-lobar tracts are massively enlarged in the wrasse and cichlid, while they are absent in the trout, the *Astyanax* surface fish and the zebrafish. 3D selective visualization of IL fiber tracts comparing the wrasse (*C. anchorago*; A), the cichlid (*N. brichardi*; B), the trout (*S. trutta*; C), the *Astyanax* surface fish (*A. mexicanus*; D), and the zebrafish (*D.* rerio; E). Lateral views are shown in (A-E), while a dorsal view of one side of the wrasse brain is shown in (F). Homologous tracts are shown in the same color across species. Besides wrasses and cichlids, no fibers connecting the pallium to the IL were detected in the other species of teleosts examined, irrespective of brain size. The main connections of the IL in these species are with the PT (blue), whereas they are with the pallium in wrasses and cichlids (ventral tract in green, dorsal tract in magenta). Local IL networks are shown in purple, and PG projections to the pallium are shown in orange. Brain regions: GR: corpus glomerulosum pars rotunda, IL: inferior lobe, PT: pretectum R: rostral; C: caudal; D: dorsal; V: ventral; L: lateral ; M: medial.

3D reconstruction of the DiI-stained fibers in the Tel and rFM of wrasses and cichlids revealed the presence of enriched fiber labeling in the Tel and rFM. Most of the IL, the ventral-most part of the rFM, exhibited high fiber density in wrasses (Figure 6A, Video 1; in purple), while fiber labeling was sparse in the other species examined (trout, Figure 6C, Video 2; *Astyanax* surface fish, Figure 6D, Video 3; zebrafish, Figure 6E, Video 4; in purple).

The telencephalic fibers in the wrasse almost completely occupied the entire pallium. These fibers converged onto the lateral forebrain bundle as they exited the telencephalon and then split again into two major tracts (Figure 6A, Video 1; in green and magenta). These connected the pallium and the structures in and around the IL (in an area which we refer to as the lobar region, Figure 4F,G), and we thus refer to these two tracts as “pallio-lobar tracts”. The ventrally located tract (Figure 6A, Video 1; in green) directly connected the pallium and the ventral IL ipsilaterally. The dorsally located tract (Figure 6A, Video 1; in magenta) coursed near the midline ipsilaterally and connected the pallium with a structure called the nucleus preglomerulosus pars commissuralis (PGc) in the lobar region, and sent minute fibers to the IL that ran through the periphery of an oval-shaped structure called the corpus glomerulosum pars rotunda (GR). The same tracts were also present in the cichlid brain, albeit more modestly, with a much smaller fiber arborization in both the pallium and IL (Figure 6B, Video 5; in green and magenta). Strikingly, in trout, zebrafish, and *Astyanax* surface fish, these tracts were not detectable, and only minimal arborization was found in the pallium and the IL (Figure 6C-E, Videos 2-4). Both PGc and GR were absent in those species, indicative of the poor development of their lobar region.

Tract tracing studies using the lipophilic dye NeuroVue, biocytin, and biotinylated dextran amine (BDA molecular weight 3000) confirmed the presence of connectivity between the pallium and the lobar region in the wrasse and cichlid brains (Figure 6-figure supplement 1, see «Materials and Methods» section). Biocytin injections in the telencephalon (Figure 6-figure supplement 1A, white asterisk) allowed us to identify the direction of the projections. Abundant labeled fibers were found in the IL (Figure 6-figure supplement 1B), while very few cell bodies were labelled (Figure 6-figure supplement 1C, white arrows). This suggests that the majority of projections were descending fibers from the pallium to the IL, with only few ascending fibers from the IL to the pallium. These fibers reached the IL through the ventral branch of the lateral forebrain bundle mentioned above.

While pallio-lobar tracts were not detectable in zebrafish with 3D reconstruction of DiI labelled fibers, biocytin injections into the dorsal telencephalon resulted in labeled fibers in the lateral forebrain bundle and terminal labeling in the IL of zebrafish. This indicates that pallial connectivity with the IL was present in this species, albeit to a lesser extent than in wrasses and cichlids.

There were two additional fiber tracts observable in all species examined. One contained projections from the sensory preglomerular complex (PG) to the pallium (Figure 6, Videos 1-5; in orange) (Bloch et al., 2020; Yamamoto & Ito, 2008), coursing rostrally and joining the lateral forebrain bundle. The most distinct branch of this tract terminated in the lateral zone of the dorsal telencephalic area (Dl) carrying visual information (Bloch et al., 2020). In the pallium of wrasses and cichlids, these visual terminals (Figure 6A,B; orange) were embedded in the arborization of the ventral pallio-lobar tract (Figure 6A,B; green).

The other tract present in all species was the one connecting the IL with the pretectum (PT) (Figure 6, Videos 1-5; in blue). In the trout, zebrafish, and *Astyanax* surface fish, this was the major tract connecting IL with the rostral aspect of the brain. In the wrasse and cichlid brain, the tract connecting the PT with IL was mediated by the GR (Figure 6A, Video 1; in blue). This only represented a small proportion of IL connectivity in those species, as the IL was also heavily connected with the pallium.

The presence of very developped pallio-lobar tracts was unrelated to absolute or relative brain size, as these tracts were not detectable in the large brained trout. Thus, the large quantity of fibers connecting the IL and the pallium in wrasses and cichlids appeared to be a remarkable feature of their brain organization compared to other species.

Overall, the presence of the pallio-lobar tracts and their extreme enlargement in wrasses may thus explain the expansion of their Tel and rFM without a corresponding increase in the relative number of cells in those structures. This increase in the relative quantity of fibers in tool-using teleost species also parallels what has been observed in the mammalian telencephalon, where primates have a larger proportion of white matter compared to rodents (Zhang and Sejnowski, 2000).

## Discussion

### Encephalization is a process of telencephalization in amniotes, but not in teleosts

Mammals and birds have taken two different trajectories of encephalization that have converged onto a process of “telencephalization”, whereby the telencephalon (and in particular the pallium) becomes massively enlarged in highly encephalized species. Our study shows that this is not the case in teleosts.

Sampling of a phylogenetically broad range of teleost species revealed that encephalization in teleosts leads to an enlargement of most of the examined brain regions, both in terms of mass and relative number of cells. That is, there was no single particularly prominent structure in highly encephalized teleosts compared to less encephalized ones. Even in the tool-using species (*C. anchorago*), the telencephalon is of a modest size, representing only 27.8% of total brain mass. This is in stark contrast to amniotes, where the telencephalon makes up 80% of total brain mass in tool-using species of primates, parrots and corvids (Herculano-Houzel et al., 2015; Olkowicz et al., 2016).

Conversely, the remarkably large structure in teleosts is rFM. In previous amniote studies (Herculano-Houzel et al., 2015; Olkowicz et al., 2016), the brain structures corresponding to rFM, TeO and rH were pooled together and called the “rest of brain” on account of their small relative size compared to the telencephalon and cerebellum. While this “rest of brain” represents merely 10 to 25% of total brain mass in primates, parrots and corvids (Herculano-Houzel et al., 2015; Olkowicz et al., 2016), it does represent 61.1% in the tool-using wrasse *C. anchorago*.

The modest telencephalon and the large “rest of brain” of teleosts, even in relatively highly encephalized tool-using species, indicates that unlike in amniotes, encephalization in teleosts is not a process of telencephalization.

### Evolution of teleost-specific brain structures with no tetrapod homolog

The rFM corresponds to the ventral part of the forebrain and midbrain, while the Tel and TeO represent the dorsal parts of the forebrain and midbrain respectively. As the rFM is large in teleosts, accounting for a quarter to a third of total brain mass, it appears that teleost brains are a lot more “ventralized” compared to amniote brains.

The lobar region (which includes the IL, GR, and PGc) in particular appeared to account for most of the rFM volume and was especially enlarged in wrasses. The IL was long considered to be of hypothalamic origin and used to be named the “inferior lobe of the hypothalamus” as a result. A recent study (Bloch et al., 2019) has demonstrated that the developmental origin of the IL is in fact mainly mesencephalic, while the cell populations surrounding the lateral recess of the hypothalamic ventricle, which represent a small part of the IL, are of hypothalamic origin. Bloch et al. (2019) has suggested that in the species with a large IL, it is mainly this mesencephalic part that becomes enlarged, and not the hypothalamic part.

GR forms the root of the lobar region in wrasses and cichlids. It is considered to have important sensory (especially visual) functions, and to project almost exclusively to the IL (Sakamoto and Ito, 1982; Shimizu et al., 1999; Yang et al., 2007). Not only does GR have no homolog in tetrapod brains, it has only evolved in some groups of teleosts (Ito and Kishida, 1975). This also appears to be the case for PGc, and we thus consider GR and PGc as specialized nuclei present only in groups of teleosts which possess large connectivity between the pallium and lobar region.

Another structure of the rFM that is also involved in sensory processing is PG, which is considered to play a role equivalent to the amniote thalamic nucleus. As our previous study has shown, it is mostly made up of cells of a mesencephalic origin (Bloch et al., 2020). In that sense, it appears that teleost brains are largely more mesencephalized than amniote brains. Altogether, sensory systems in teleosts and tetrapods are not as conserved as previously thought but have evolved independently in each lineage.

Teleosts thus display marked differences in the organization of their brains compared to amniotes, with mesencephalic structures accounting for a much larger proportion of total brain mass and playing a prominent role in sensory processing.

### Different ways of evolving tool-using brains

Our current study revealed that in the wrasse and cichlid brains, the IL is highly connected with the pallium. This seems to be especially apparent in tool-using species. As some previous studies already suggested (Bloch et al., 2019; Schmidt, 2020), this challenges the previous notion that the IL is merely a food motivation center (Demski, 1973; Demski and Knigge, 1971; Muto et al., 2017; Roberts and Savage, 1978). IL receives gustatory information (Morita et al., 1980, 1983; Rink and Wullimann, 1998; Yang et al., 2007), and due to its position directly next to the hypothalamus, it was thought to be homologous to the lateral hypothalamus of mammals (Roberts and Savage, 1978). Direct electrical stimulation of the IL resulted in behaviors such as bitting at a mirror or snapping at objects in freely moving fish (Demski, 1973; Demski and Knigge, 1971), and IL activation was found during detection of moving objects in larval zebrafish (Muto et al., 2017). With the assumption that the IL was homologous to the mammalian hypothalamus, these functional data have been interpreted as the IL playing a role in feeding behaviors. However, since this previous view of IL homology has been shown to be erroneous (Bloch et al., 2019), a reinterpretation of this data is necessary.

In addition to gustatory inputs, IL receives visual inputs from the TeO via the pretectum (PT) (Muto et al., 2017; Sakamoto and Ito, 1982; Shimizu et al., 1999; Yang et al., 2007). In the species where GR is present, it has been suggested that IL also receives auditory (Sakamoto and Ito, 1982) and somatosensory (Shimizu et al., 1999) information. As a result, IL has also been proposed to be a multi-sensory integration center (Ahrens and Wullimann, 2002; Rink and Wullimann, 1998; Shimizu et al., 1999; Yang et al., 2007). In addition, since its main output is to the lateral valvular nucleus, which projects to the cerebellum (Yang et al., 2004, 2007), its functions may be motor-related. This sensory input and motor output connectivity pattern in IL is rather similar to what has been found in the amniote pallium. As the teleost pallium itself receives sensory information of different modalities (e.g. auditory and visual inputs via PG), the IL seems to serve as another sensory integration center in the teleost brain (Figure 7).

**Figure 7.**
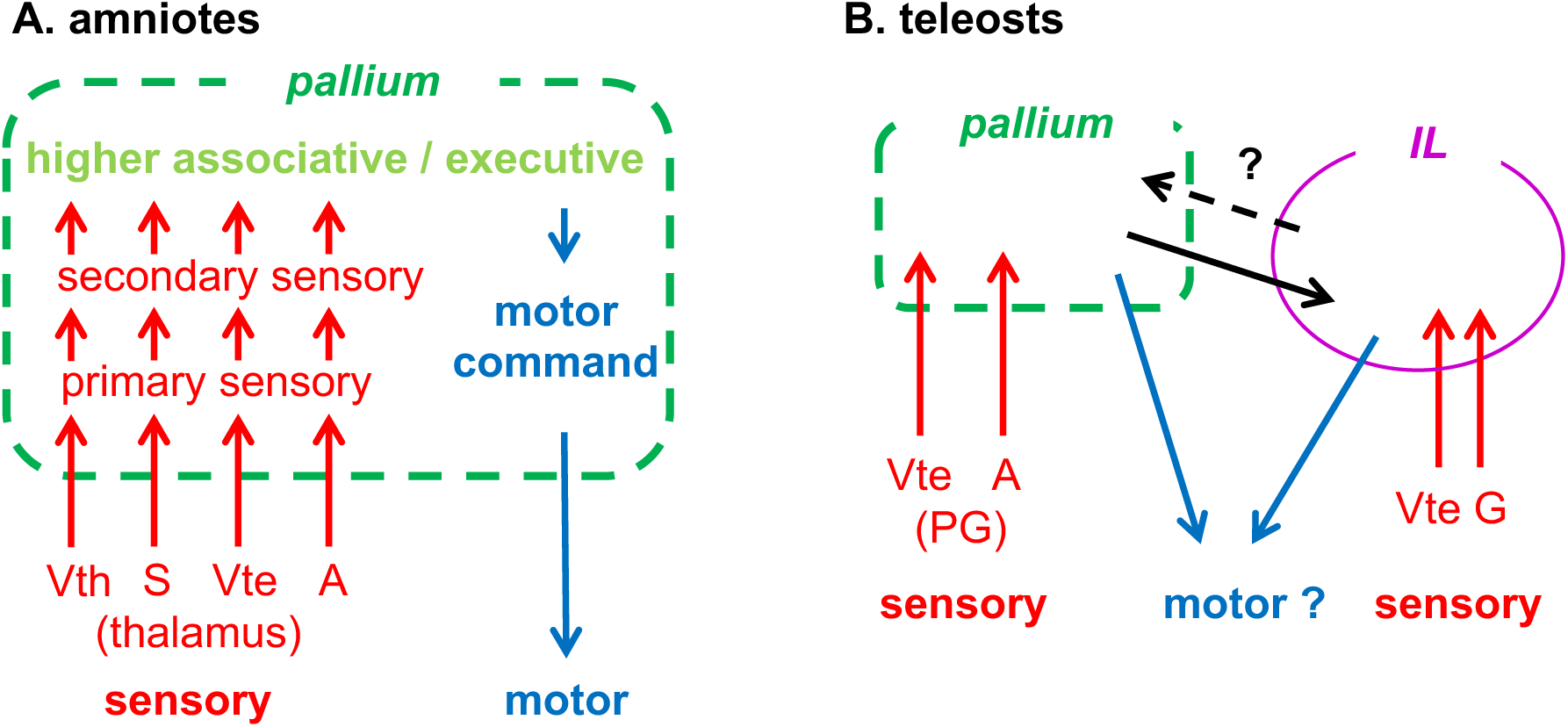
Comparison of functional connectivity in relation to sensory inputs and motor outputs in amniotes and teleosts. (A) Simplified diagram showing input/output connectivity of the pallium commonly found in mammals and birds (analogous, not necessarily homologous). Sensory inputs are shown in red, while motor outputs are shown in blue. The primary sensory areas in the pallium receive modal-specific sensory inputs from subtelencephalic sensory nuclei, mainly through the thalamus in the case of tetrapods. Note that there are two major visual pathways terminating in the pallium both in mammals and birds. The diagram is modified from Yamamoto and Bloch (2017). (B) Simplified diagram showing input/output connectivity of the pallium and IL in teleosts. The sensory afferents to the pallium in teleosts are mainly mediated via the PG instead of the thalamus. In addition to the pallium, IL receives sensory inputs of different modalities, here showing only visual and gustatory, which are the dominant ones. The pallium and IL are highly connected in some teleost groups such as wrasses and cichlids. Sensory modalites: A: auditory, G: gustatory, S: somatosensory, Vte: visual (tectofugal), Vth: visual (thalamofugal).

The presence of multimodal inputs to the IL is likely to be a common feature in teleosts, but the particularity of the wrasse and cichlid brains is the IL’s intense connectivity with the pallium. The major connectivity of the IL of other fish like trout, *Astyanax* surface fish, and zebrafish is with the PT, which is involved in stereotyped movements such as the optokinetic response (Kubo et al., 2014; Portugues et al., 2014) or the prey detection J-turn in larval zebrafish (Fajardo et al., 2013). Those types of movements are sufficient for simple foraging behaviors without flexibility. It is then possible that the elaborated connectivity with the pallium present in wrasses and cichlids may have allowed for the emergence of their complex behavioral repertoire. The large gustatory inputs of the IL may for instance be involved in different functions than simply eating in these species. As fish do not have hands, they use their mouth to manipulate objects, and could likely have fine discriminative touch and motor control via the lips and oral cavity, functions which could involve the IL. Apart from tool use in wrasses, cichlids display object play and elaborate nest building behaviors (Burghardt et al., 2015; Johnson et al., 2020; Paśko, 2010; York et al., 2015), which also require this kind of precise motor control.

The presence of a higher-order association center in the teleost pallium has hardly been investigated so far. In mammals and birds, the sensory association areas are located in the periphery of the primary sensory areas (Atoji and Wild, 2012; Goodale and Milner, 1992; Güntürkün, 2005; Jarvis et al., 2013; Stacho et al., 2016; Swenson and Gulledge, 2017). If the teleost association areas are organized in the same manner, the area where the arborization of the ventral pallio-lobar tract is located (Figure 6A,B; green) would be a good candidate for the visual association area. The Dl, the putative teleost primary visual area (Figure 6A,B; orange pallial arborizations), projects to the surrounding pallial areas including the central part of the pallium (Hagio and Yamamoto, 2023; Yamamoto and Ito, 2008), which in turn project to the IL. These observations raise the possibility that IL may also be a part of the higher-order areas of the teleost brain. Additional connectivity and functional studies are required to verify how higher-order areas are organized in the teleost brain.

In conclusion, our findings revealed that the encephalization process in teleosts is different from what has previously been described in amniotes. While the pallium also appears to be important for higher-order cognitive functions in teleosts, the large pallio-lobar tracts in the tool-using fishes demonstrate the functional importance of the IL in relation to the pallium, which may be critical for such complex behaviors. Since the IL has no homolog in amniotes, at least three different brain organizations enabling higher-order cognitive functions may have evolved independently in mammals, birds and teleosts.

## Materials and Methods

All animal experiments were conducted in compliance with the official regulatory standards of the French Government and in compliance with the official Japanese regulations for research on animal, and the regulations on Animal Experiments in Nagoya University.

### Study animals and brain sampling

11 species of teleost were examined: a group of 3 wrasse species (*Choerodon anchorago*, *Labroides dimidiatus, Thalassoma hardwicke*), for which complex behaviors (tool use and social cognition) have been reported, 4 cichlid species (*Maylandia zebra*, *Neolamprologus brichardi*, *Ophthalmotilapia boops*, *Amatitlania nigrofasciata*), which are phylogenetically close to wrasses and are capable to a lesser extent of complex behaviors, and a group of 4 other species (the medaka (*Oryzias latipes*), zebrafish (*Danio rerio*), *Astyanax* surface fish (*Astyanax mexicanus*), and trout (*Salmo trutta*)) for which no such behavior has been observed.

Adult individuals of zebrafish (*Danio rerio*), medaka (*Oryzia latipes*) and *Astyanax mexicanus* were obtained from the animal facility in NeuroPSI (Saclay, France). Adult trouts (*Salmo trutta*) were sourced from the animal facility at INRAE (Jouy-en-Josas, France). *Neolamprologus brichardi*, *Amatitlania nigrofasciata* and *Danio rerio* individuals used for tract-tracing with BDA and biocytin were obtained from local dealers in Japan.

Sexually mature individuals of both sexes of wrasse and cichlid species were sourced from commercial providers (*Choerodon anchorago*, *Labroides dimidiatus*, *Thalassoma hardwicke*: Marine Life (Paris, France); *Maylandia zebra* and *Ophthalmotilapia boops*: Abysses (Boissy-Saint-Léger, France); *Amatitlania nigrofasciata*: Abysses, Aquariofil.com (Nîmes, France), *Neolamprologus brichardi*: Abysses, Aquariofil.com). Wrasses were wild caught and tended to be young adults, but one large adult of *Choerodon anchorago* weighing around ten times as much as the other individuals was also sampled. Statistical analysis showed that the data from this large individual did not impact the statistical significance of our results (see Supplementary file 1).

Zebrafish and medaka specimens were euthanized in ice-cold water, weighed on a precision scale and fixed in ice-cold 4% paraformaldehyde (PFA; Electron Microscopy Science) in 0.01M phosphated buffer saline containing 0.1% Tween 20 (PBST; Fisher Scientific). All other fish specimens were euthanized by an overdose of 0.4% tricaine methanesulfonate added to fish water (MS222; Sigma-Aldrich), weighed, and immediately perfused transcardially with 4% PFA in PBS. 24 hours post-fixation, brains were dissected, weighed on a precision scale, and kept in 4% PFA in PBS for another 24 hours before being transferred in anti-freeze solution (30% glycerol, 30% ethylene glycol, 30% H20, 10% PBS 10X) and kept at −20°C for later use. Brains used for NeuroVue tract-tracing were kept in 4% PFA at 4°C until use.

### Isotropic fractionator

#### The medaka brains (n=5) were left undissected

The brains of n=5 individuals of each species, except the trout (n=4), *M. zebra* (n=3), *C. anchorago* (n=4), *L. dimidiatus* (n=3), *T. hardwicke* (n=3) and *O. boops* (n=3) were rinsed in PBS and embedded in 3% agarose containing 1% Tween 20 and sectioned at 300 µm in the frontal plane with a vibratome (Leica VT 1200S). Under a stereomicroscope (Olympus SZX7), the brain was manually dissected using a microsurgical knife (Stab Knife 5mm, 15 degrees; Surgical Specialties Corporation) into five regions following the rostro-caudal and dorso-ventral axis (Figure 4).

The dorsal part of the secondary prosencephalon, which includes the telencephalon and the dorso-rostral part of the optic recess region (ORR) (Yamamoto et al., 2017; Yamamoto and Bloch, 2017), was excised. This region was labelled “telencephalon” (Tel). The second region dissected was the dorsal part of the mesencephalon, which includes the tectum opticum and the torus semicircularis and was labelled “optic tectum” (TeO). The third region included the ventral part of the secondary prosencephalon (i.e., the hypothalamus), the diencephalon and the ventral part of the mesencephalon (i.e., the tegmentum and the inferior lobe) and was labelled “rest of the forebrain/midbrain” (rFM). The fourth excised region was the dorsal part of the rhombencephalon (i.e., the cerebellum) (Cb). Finally, all the other hindbrain structures, including the medulla oblongata, were labelled “rest of the hindbrain” (rH). Sections were dried with a paper towel, weighed on a precision scale and kept in 4% PFA for later use.

The number of cells in the five main regions of the teleost brain was determined using the isotropic fractionator method (Herculano-Houzel and Lent, 2005). This method produces results similar to unbiased stereology (Bahney and von Bartheld, 2014; Miller et al., 2014). Each structure was manually homogenized in 40 mM sodium citrate (Sigma-Aldrich) with 1% Triton X-100 using a Tenbroeck tissue grinder (Ningbo Ja-Hely Technology Co., Ningbo, China). Once an isotropic suspension of isolated cell nuclei was obtained, the suspension was then centrifuged, and the supernatant was collected. The cell nuclei in both the suspension pellet and the supernatant were stained by adding PBS with 1% diamino-phenyl-indol (DAPI; Sigma-Aldrich).

Additionally, a predetermined volume of PBS was added to the suspension to adjust the nuclei density for counting.

To determine the total number of cells in the tissue, four 10 µL aliquots of the suspension and of the supernatant were counted under an epifluorescence microscope (Axio Imager, Zeiss) with X200 magnification using a Blaubrand Malassez counting chamber (Brand Gmbh, Wertheim, Germany). Mean nuclear density in the suspension and the supernatant was multiplied by their total volume and added up to determine the total number of cells in the brain tissue.

To determine the total number of neurons in each sample, we initially aimed at performing an anti-NeuN immunoreaction in PBS using anti-NeuN antibodies. However, after testing multiple antibodies (anti-NeuN rabbit Antibody, ABN78 & ABN78C3, Merck; anti-NeuN rabbit Antibody, ab177487, Abcam; anti-NeuN mouse Antibody, MAB377, Merck) and increasing antibody concentrations (up to 1:50), we were unable to obtain reliable neuronal nuclear staining. Further tests on brain sections also failed to label teleost neuronal nuclei with NeuN in a consistent manner, suggesting that this tool was not appropriate for teleost tissues. Consequently, we decided to present data on total cell numbers for our brain samples.

### Whole-brain clearing and staining

Lipophilic dye was applied to n=2 whole brains of *D. rerio*, *A. mexicanus*, *N. brichardi*, *C. anchorago* and *S. trutta* for fiber bundles tracing.

Brains stored in anti-freeze solution at −20°C were washed with PBST for at least 1 day. Samples were bleached for 2 hours under intense lighting (>10000 lux, GVL-SPOT-50-FIXV4-230VAC, GreenVisuaLED) in a fresh depigmentation solution of 5% H_2_O_2_, 0.05% sodium azide in PBS. The bleached samples were thoroughly washed with PBST overnight and were then subjected to a size-dependent delipidation step in CUBIC-L (Tainaka et al., 2018). They were first immersed in a mixture of 50% PBST/50% CUBIC-L overnight under gentle shaking followed by an incubation in CUBIC-L at 37°C under agitation for 1-2 days for *D. rerio*, 3 days for *A. mexicanus*, 4 days for *N. brichardi* and 6 days for *C. anchorago* and *S. trutta* with solution renewed once. Delipidated specimens were washed with PBST for at least 4 hours prior to staining.

Staining was performed in solutions that were originally designed for immunostaining of zebrafish larvae (Lempereur et al., 2022). Samples were immersed in a blocking solution of 10% normal goat serum, 10% DMSO, 5% 1M PBS-glycine, 0.5% Triton X-100, 0.1% sodium deoxycholate, 0.1% IGEPAL CA-630 and 0.1% saponin in PBST overnight at 37°C under gentle shaking. Subsequently, specimens were stained with 2 µg/ml of DiI (D282; ThermoFisher Scientific) in a solution of 2% NGS, 20% DMSO, 0.05% sodium azide, 0.2% Triton-X100, 10 µg/mL heparin and 0.1% saponin at 37°C under rotation for a specimen-dependent duration.

After a last washing step in PBST, refractive index (RI) matching was carried out in weakly basic CUBIC-R solution (Tainaka et al., 2018). Brains were soaked in a mixture of 50% PBST/50% CUBIC-R overnight under agitation and then kept in CUBIC-R (RI = 1.52) prior to mounting.

### Whole-brain 3D imaging

RI matched samples were embedded in a filtered (pore size 5.0 µm) melted agarose solution containing 2% agarose, 70% CUBIC-R in distilled H_2_O. CUBIC/agarose gels were immersed in CUBIC-R at RT for a minimum of 1 day to homogenize RIs.

Images were acquired with two commercial light-sheet fluorescence microscopes. Acquisitions were performed with an Ultramicroscope II (Miltenyi Biotec) using a 1.1x NA 0.1 MI PLAN objective and a DC57 WD17 0 dipping cap coupled to a 2x magnification lens, or a LVMI-Fluor 4x/0.3 WD6 objective without additional magnification. A Lightsheet 7 (Zeiss) equipped with 10x 0.2 foc illumination and 5x 0.16 foc detection optics was also used. According to brain size and the type of microscope images were acquired from dorsal or sagittal view. Cotton seed oil was poured on the surface of the imaging medium as an impermeable layer to avoid evaporation-induced RI changes during imaging. 16-bit images were acquired by a pco.edge 5.5 sCMOS camera (2560 x 2160 pixels, pixel size 6.5 µm x 6.5 µm) on the Ultramicroscope II or a pco.edge 4.2 sCMOS camera (1920 × 1920 pixels, pixel size 6.5 µm × 6.5 µm) on the Lightsheet 7, following sample excitation with laser 488 and 561 nm. The z-step size was fixed to 6 µm on the Ultramicroscope II and 5.176 µm on the Lightsheet 7, which represents nearly half of the theoretical lightsheet thickness.

### 3D image reconstruction and manual segmentation

For the inter-species comparison of the anatomy of the tracts connecting the Tel with the rFM, these structures were segmented manually using the 3D visualization and reconstruction software Amira 2019 (Thermo Fisher Scientific).

The combination of the overall size of the specimens and the required resolution/voxel size demanded tiled image acquisition. The resulting image stacks were merged using the Grid/Collection stitching plugin (Preibisch et al., 2009) in Fiji (Schindelin et al., 2012).

In preparation for the manual segmentation, the signal-to-noise ratio of the merged data was improved by subtracting the gaussian noise (Fiji, Gaussian Blur 3D, Kernel 10,10,10) from the original data. After manual segmentation of the original data and the denoised data by an unbiased researcher, the defined regions were refined by multiplying the denoised data with the individual binary masks of the segmentations.

The 3D reconstructions in Figure 6 were produced on n=2 brains for each species by selective visualization of the denoised features under investigation in this study within the framework of the overall anatomy of the corresponding brains.

### Tract-tracing with NeuroVue

In order to confirm the presence of the IL fiber tracts visualized with DiI staining, tract-tracing experiments were performed using NeuroVue (Polysciences), a lipophilic dye which allows both retrograde and anterograde tracing and can be used on fixed brain tissue (Duncan et al., 2011). Small triangular pieces of NeuroVue filter paper were inserted into the IL of n=3 specimens of *A. mexicanus*, *N. brichardi* and *C. anchorago*, and into the pallium of n=3 specimens of *N. brichardi*. Brains were then incubated at 36°C in 4% PFA in PBS for 4 (*A. mexicanus*) to 12 days (*C. anchorago*).

Following incubation, 80 µm sections were cut with a vibratome (Leica VT1200S) in the sagittal plane for *A. mexicanus*, and for the *N. brichardi* specimens which were injected into the telencephalon. IL-injected *N. brichardi* and *C. anchorago* specimens were cut in the frontal plane. Sections were then treated with DAPI before being mounted on glass slides with VectaShield mounting medium (Vector Laboratories). Sections were imaged using a Leica SP8 confocal microscope.

### Tract-tracing with BDA and biocytin

BDA (molecular weight 3000; Thermo Fisher Scientific Molecular Probes) and biocytin (Sigma-Aldrich) was injected *in vivo* into the pallium of two species of cichlids of both sexes, *N. brichardi* (n = 5; standard length: 30-49 mm), *A. nigrofasciata* (n = 5; standard length: 45-55 mm) and zebrafish *D. rerio* (n=2; standard length: 30 and 35 mm). Fish were anesthetized by immersion in water containing 150-180 mg/L MS222 and set in a device for physical restraint. Water containing 70-80 mg/L MS222 was perfused through the gill for aeration and to maintain the anesthetic condition. A dorsal portion of the cranium was opened to expose the brain. For BDA injections, a glass microelectrode (tip diameters: 12-16 µm) filled with 0.75% BDA solution in 0.05M Tris-HCl-buffered saline (TBS; pH 7.4) was driven into the pallium with a manipulator (MN-3; Narishige). BDA was injected iontophoretically with square current pulses (+5 µA, 0.5 Hz, 50% duty cycle) passed through the electrodes at three to six places of the pallium each for 5 minutes with a stimulator (SEN-3301; Nihon Kohden, Japan). For biocytin injections, crystals of biocytin were inserted with a minute insect pin into three to six places of the pallium. After the injection, the cranial opening was closed with either plastic wrap (small fish) or dental cement (Ostron II; GC Dental Products, Japan). Postoperative fish were maintained in aquaria for 21-30 hours. The fish were then deeply anesthetized with MS222 (over 200 mg/L) and perfused through the heart with 2% PFA and 1% glutaraldehyde in 0.1 M phosphate buffer (PB), pH 7.4. The brains were removed from the skull and post-fixed in the same fixative at 4°C for 6-8 hours.

The fixed brains were cryo-protected by immersion in 0.1 M PB containing 20% sucrose at 4°C. Cryo-protected brains were embedded in 5% agarose (type IX, ultra-low gelling temperature; Sigma-Aldrich) containing 20% sucrose and frozen in n-hexane at −60°C. Then, frontal sections were cut at a thickness of 40 µm on a cryostat and mounted on gelatin-coated glass slides. The sections were dried and washed once with 0.05 M TBS containing 0.1% Tween 20 (TBST) and twice with TBS. To quench non-specific peroxidase activities, sections were steeped in methanol containing 0.3% H_2_O_2_ and washed three times with TBS and once with 0.03% TBST. Sections were then incubated with a solution of avidin-biotin-peroxidase complex (1:100; VECTASTAIN Elite ABC Standard Kit, Vector Laboratories) overnight. Afterwards, sections were incubated for one hour with 0.05% 3,3’-diaminobenzidine (Sigma-Aldrich) solution in 0.1M PB containing 0.04% nickel ammonium sulfate and 0.01% H_2_O_2_. The reaction was stopped by washing four times with TBS, and the sections were counterstained with 0.05-0.1% cresyl violet, dehydrated, and coverslipped with Permount (Fisher Scientific).

### Statistics

To determine whether brain mass, body mass, and total number of cells in the brain are correlated in teleosts, a nonparametric Spearman rank correlation test was used on log-transformed data. Previously published data on birds (Olkowicz et al., 2016) and mammals (Herculano-Houzel et al., 2015) were used for comparison. If a P<0.05 value was found, reduced major axis (RMA) regressions were calculated using the SMATR package (Warton et al., 2012) in RStudio. To compare scaling among taxonomic groups, an analysis of covariance (ANCOVA) with post-hoc Sidak corrected pairwise comparisons was used to check for significant differences in the slopes of the regression lines. In groups for which the slopes were statistically homogeneous, the regression lines were compared based on the differences in their intercepts.

In order to determine the degree of encephalization of the teleost species sampled in this study, a phylogenetically corrected brain-body allometric slope was estimated using phylogenetically generalized least squares regression test (PGLS) at the Class level on species means of log^10^ brain and log^10^ body mass data of the species sampled in this study along with previously published actynopterygian data by Tsuboi et al. (2018) (Tsuboi et al., 2018) using RStudio with the CAPER package. Residual variance was modelled according to Brownian motion (Felsenstein, 1985) and phylogenetic signal was estimated using Pagel’s λ (Lynch, 1991). Phylogenetic relationships between teleost species were based on previously published phylogenetic trees (Rabosky et al., 2013). Encephalization was then determined by extracting the residuals of log^10^-log^10^ brain and body mass for each species of the dataset to remove allometry in brain size (Sol et al., 2016). The 11 species studied were ranked based on the value of their residual (Table 2).

To determine whether there exists a correlation between the degree of encephalization and relative mass and relative number of cells (expressed as the percentage of total brain mass and percentage of total brain cells, respectively) of the five major brain structures dissected, a nonparametric Spearman rank correlation test on species means was used, as there was no way to ascertain the normal distribution of these data. Nonparametric Kruskal-Wallis and Dunn’s post hoc tests were used to assess the inter-species differences in relative mass, absolute and relative number of cells in the five dissected brain structures, while nonparametric Mann-Whitney tests were used to compare the relative mass and relative number of cells in major brain structures between groups as normality was not verified for all species or groups. All tests were performed in GraphPad Prism. Statistical parameters are given in the Figure legends.

## Acknowledgements

We thank Jean-Michel Hermel and Naomie Pradère (NeuroPSI, CNRS/Université Paris-Saclay) for their help with brain sampling. We thank Dimitri Rigaudeau (INRAE, Jouy-en-Josas) for providing us trouts and zebrafish, the DECA team (NeuroPSI) for *Astyanax* specimens, Joël Attia (Université de Saint-Etienne) for cichlids, Anthony Herrel (Muséum National d’Histoire Naturelle, Paris) for wrasses, as well as the members of the animal facility (CNRS UMS2010, INRA UMS1451, Université Paris-Saclay), especially Krystel Saroul and Christophe de Medeiros for fish care. We thank members of NeuroPSI for technical support, and the MIMA2 platform (https://doi.org/10.15454/1.5572348210007727E12; INRAE, Jouy-en-Josas) for access to their lightsheet microscope. Finally, we thank Florian Razy-Krajka and Rose Tatarsky for their help in improving the manuscript.

---

**Figure 2-figure supplement 1.**
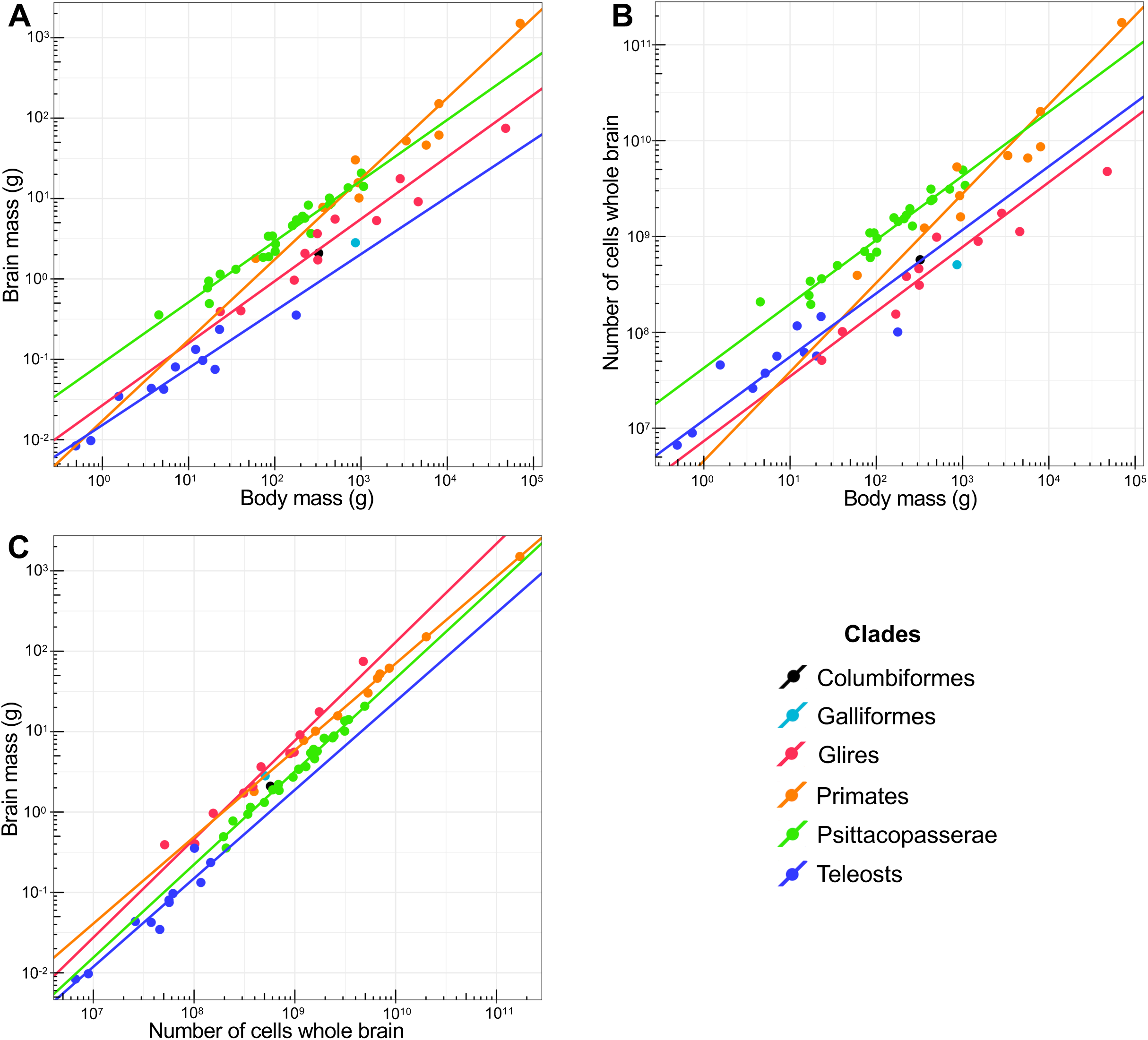
Teleosts have small, cell-dense brains that contain more cells than the brains of rodents of similar body mass even after removal of the large *Choerodon anchorago* individual from the analysis. (A-C) The fitted reduced major axis (RMA) regression lines are displayed only for correlations that are significant. Each point represents the mean value of a species. X and y axes are in log^10^ scales. All regression lines are significantly different, except for the regression lines of Glires and Primates in plot (C). (A) Brain mass plotted as a function of body mass. Teleosts have smaller brains than birds and mammals of similar body mass. (B) Total number of cells in the brain plotted as a function of body mass. Teleost brains contain less cells than bird and primate brains, but more cells than the brains of rodents of similar body mass. (C) Brain mass plotted as a function of total number of cells in the brain. Cellular density inside the teleost brain is higher than in birds and mammals. See also Table 1. For statistics, see Supplementary file 1. Glires = rodents and lagomorphs. Psittacopasserae = Passeriformes (songbirds) and Psittaciformes (parrots).

**Figure 3-figure supplement 1.**
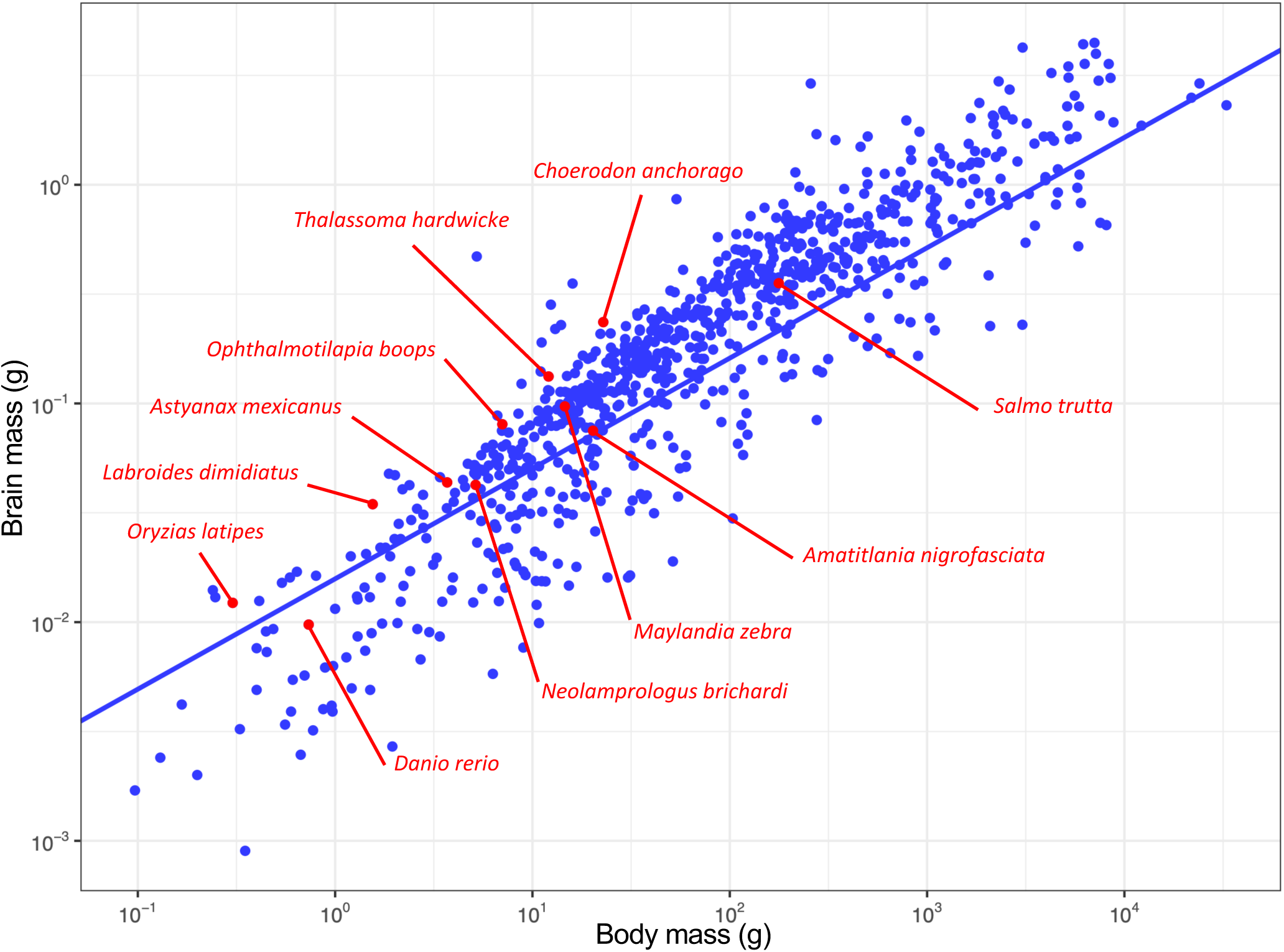
Encephalization in 11 species of teleosts compared to a large dataset of actinopterygians after removal of the large *Choerodon anchorago* individual. Brain mass is plotted as a function of body mass, and the phylogenetically corrected (phylogenetically generalized least squares regression test, PGLS) allometric line is shown. Each point represents the mean value of a species. X and y axes are in log^10^ scales. The phylogenetic regression slope for actinopterygians is of 0.50 ± 0.01. Adjusted R^2^: 0.8379, t=65.891, p<0.0001. See also Table 2 and Supplementary file 1.

**Figure 3-figure supplement 2.**
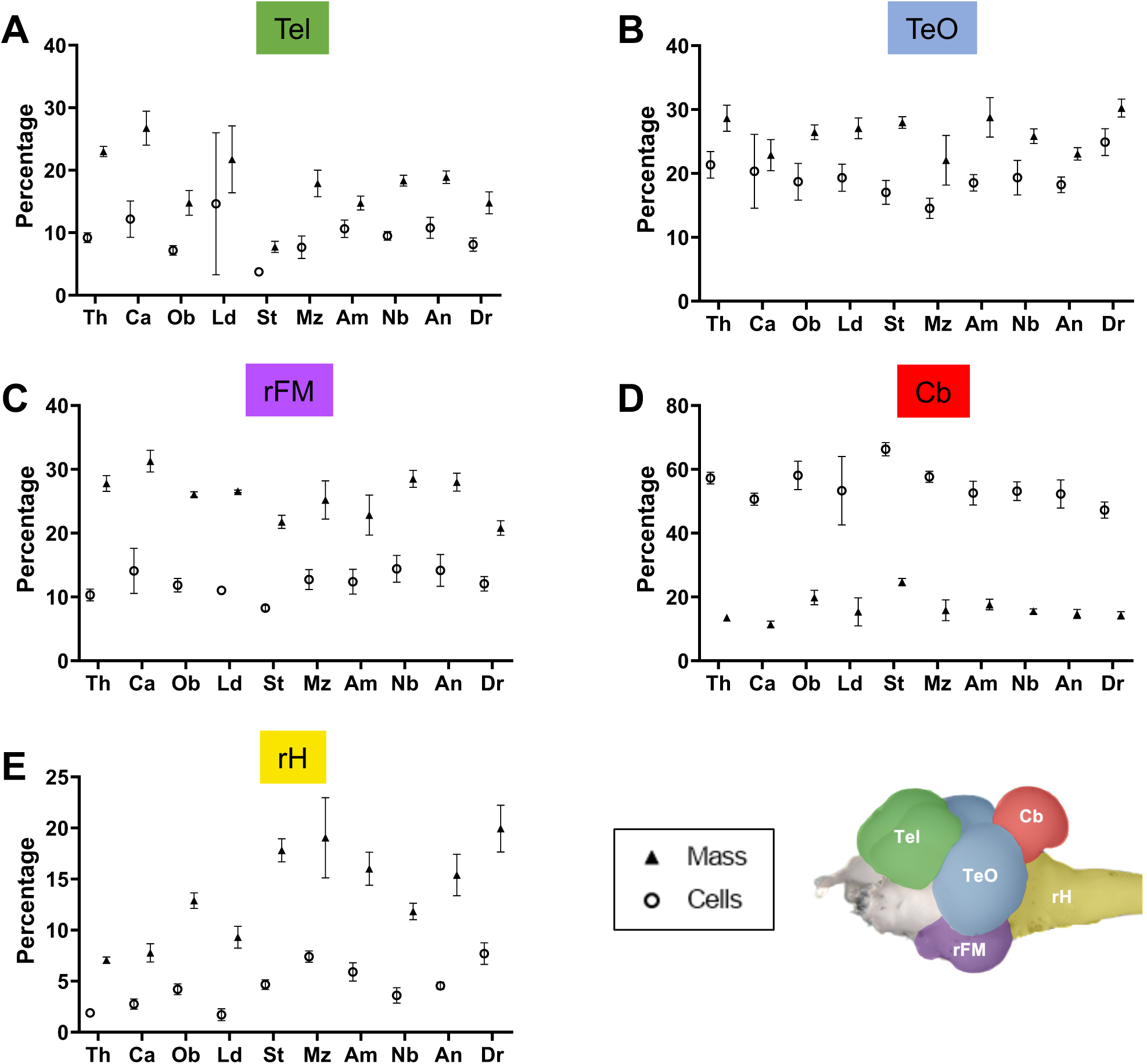
Encephalization in teleosts is not correlated with an increase in the relative mass or relative number of cells in the telencephalon (Tel). (A-E) Relative mass (triangles) and relative number of cells (open circles) of the Tel (A), TeO (B), rFM (C), Cb (D), and rH (E) of ten species of teleosts. Species are ranked from most encephalized (left) to least encephalized (right). Each point represents the mean value of a species. Error bars show SD. Spearman’s rank correlation test was used. Relative mass and relative number of cells in the rH (E) are negatively correlated with encephalization (Spearman r: -0.709, p=0.027 and Spearman r: -0.673, p=0.039), whereas no significant correlation exists for the other regions. Species: Am: *Astyanax* mexicanus; An: *Amatitlania nigrofasciata*; Ca: *Choerodon anchorago*; Dr: *Danio rerio*; Ld: *Labroides dimidiatus*; Mz: *Maylandia zebra*; Nb: *Neolamprologus brichardi*; Ob: *Ophtalmotilapia boops*; St: *Salmo* trutta; Th: *Thalassoma hardwicke*; Brain regions: Cb: cerebellum, rFM: rest of the forebrain/midbrain; rH: rest of the hindrain; Tel: telencephalon; TeO: optic tectum.

**Figure 3-figure supplement 3.**
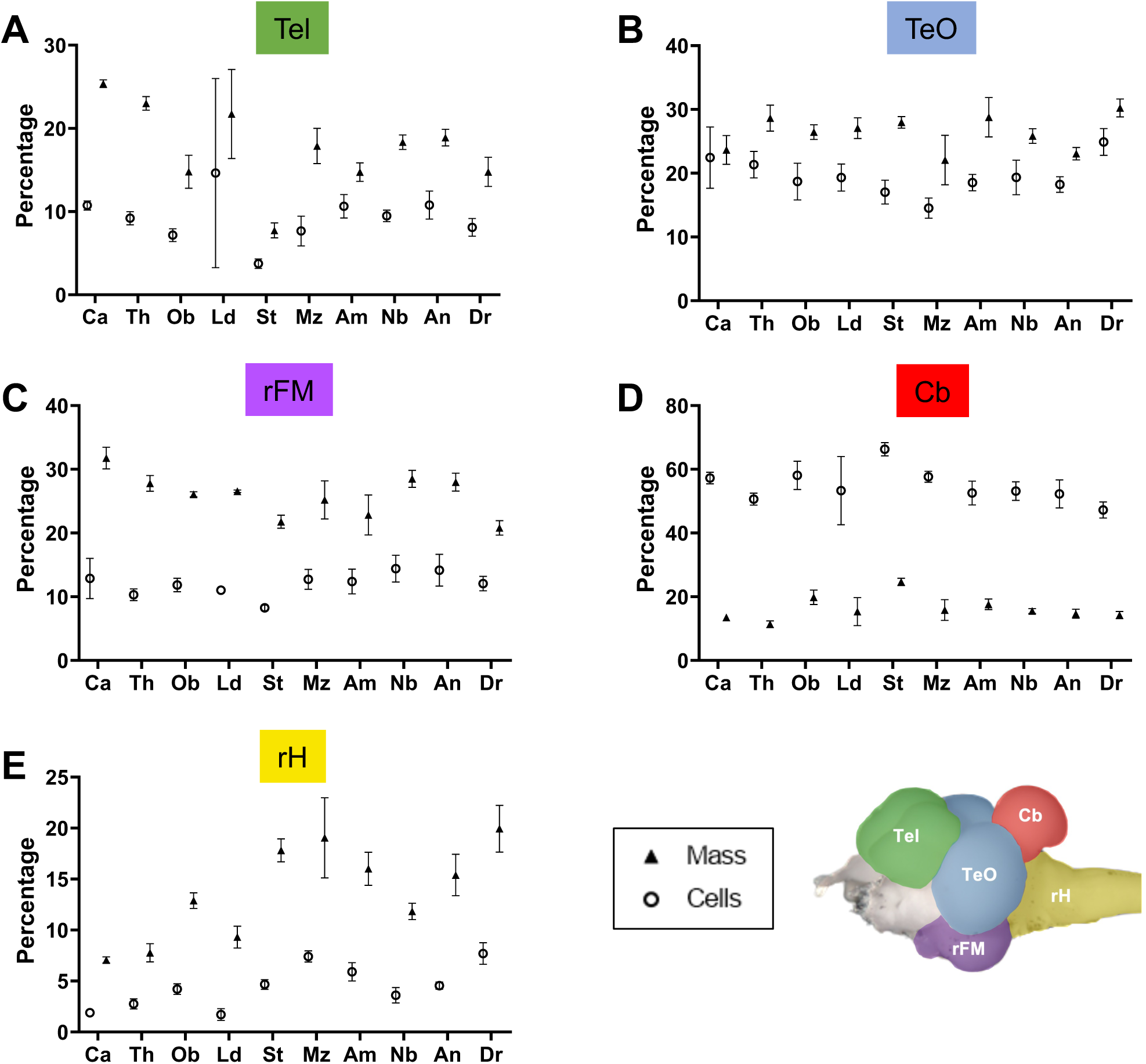
Encephalization in teleosts is not correlated with an increase in the relative mass or relative number of cells in the telencephalon even after removal of the large *Choerodon anchorago* individual from the analysis. (A-E) Relative mass (triangles) and relative number of cells (open circles) of the Tel (A), TeO (B), rFM (C), Cb (D), and rH (E) of ten species of teleosts. Species are ranked from most encephalized (left) to least encephalized (right). Each point represents the mean value of a species. Error bars show SD. Spearman’s rank correlation test was used. Relative mass and relative number of cells in the rH (E) are negatively correlated with encephalization (Spearman r: -0.697, p=0.031 and Spearman r: -0.661, p=0.044), whereas no significant correlation exists for the other regions. Species: Am: *Astyanax* mexicanus; An: *Amatitlania nigrofasciata*; Ca: *Choerodon anchorago*; Dr: *Danio rerio*; Ld: *Labroides dimidiatus*; Mz: *Maylandia zebra*; Nb: *Neolamprologus brichardi*; Ob: *Ophtalmotilapia boops*; St: *Salmo* trutta; Th: *Thalassoma hardwicke*; Brain regions: Cb: cerebellum, rFM: rest of the forebrain/midbrain; rH: rest of the hindrain; Tel: telencephalon; TeO: optic tectum. See also Supplementary file 1.

**Figure 5-figure supplement 1.**
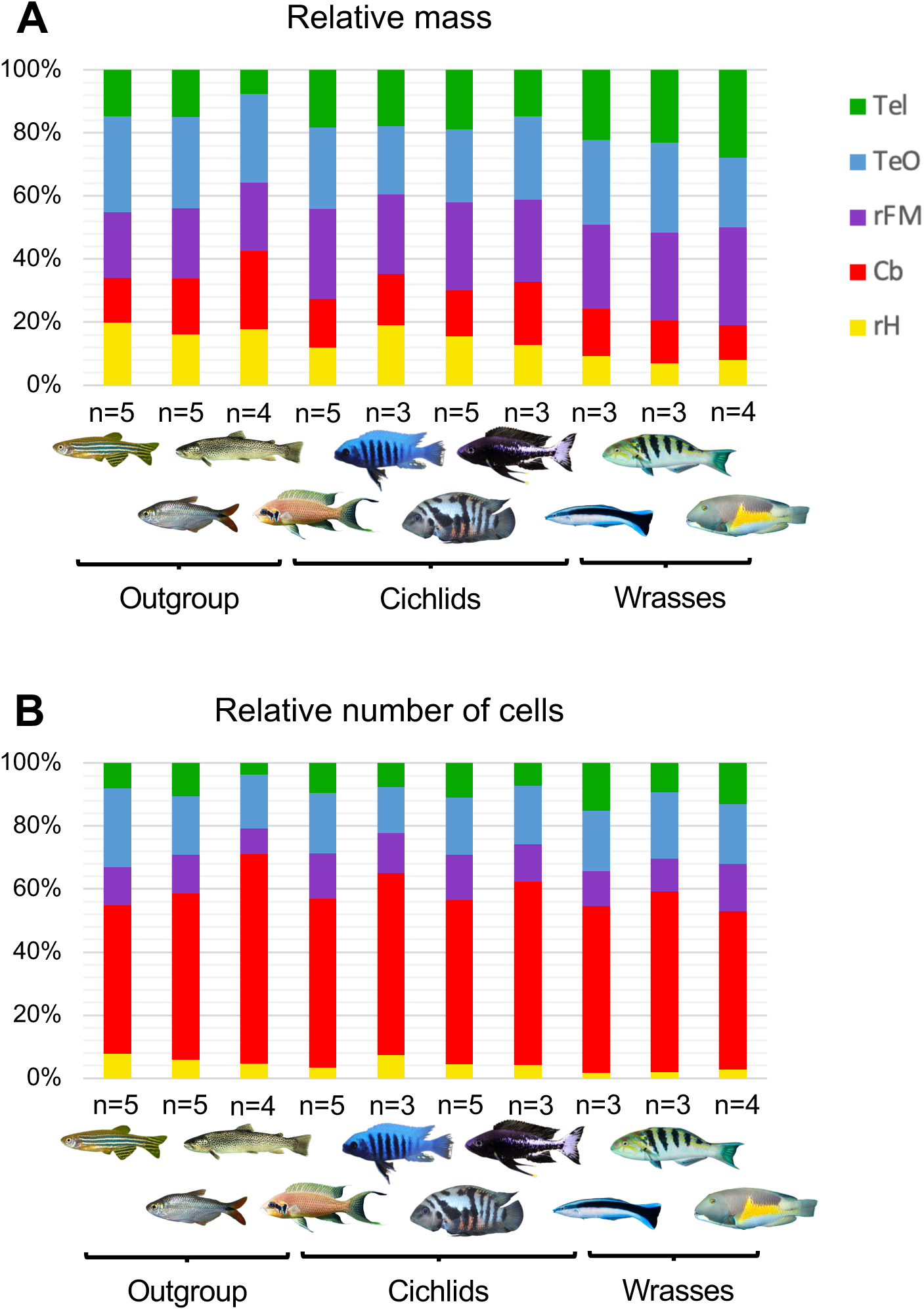
Comparison of relative mass and number of cells in the brains of teleosts. Mass distribution and cellular composition of teleost brains appear to be similar across phylogeny when species are compared one to one. Relative mass (A) and number of cells (B) of the Tel, TeO, rFM, Cb and rH of ten teleost species. Values are mean percentages per species. Significant differences were found in the absolute and relative number of cells, as well as in the relative mass in all five structures (Kruskal-Wallis, p<0.001, p<0.05 and p<0.05 in all cases, respectively). However, post-hoc pairwise comparisons revealed significant differences that were inconsistent across species and brain structures, the only consistently found difference being between *D. rerio* and *C. anchorago* in the absolute number of cells across all structures (Dunn’s test, p<0.05 in all cases), as well as in the relative mass of the rH between *A. mexicanus*, *C. anchorago* and *T. hardwicke* (Dunn’s test, p=0.0307 and p=0.0317, respectively), and in the relative mass of the Tel between *C. anchorago* and *S. trutta* (Dunn’s test, p=0.0254). Species from left to right: “outgroup” (*Danio rerio*, *Astyanax mexicanus*, *Salmo trutta*), Cichlids (*Neolamprologus brichardi*, *Maylandia zebra*, *Amatitlania nigrofasciata*, *Ophthalmotilapia boops*), Wrasses (*Labroides dimidiatus*, *Thalassoma hardwicke*, *Choerodon anchorago)*. Brain regions: Cb: cerebellum, rFM: rest of the forebrain/midbrain; rH: rest of the hindrain; Tel: telencephalon; TeO: optic tectum.

**Figure 5-figure supplement 2.**
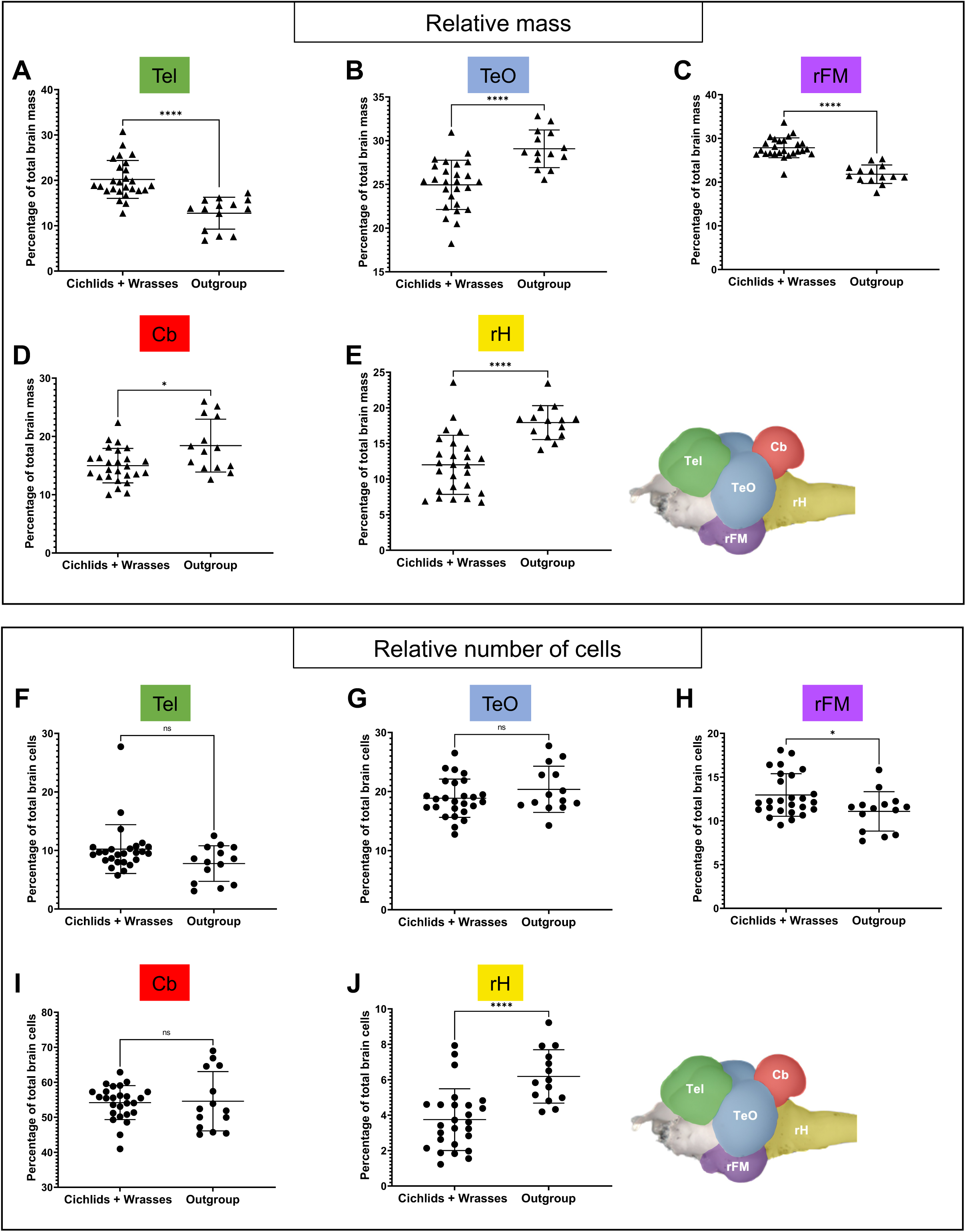
Wrasses and cichlids have a relatively larger telencephalon (Tel) and rest of the forebrain/midbrain (rFM) compared to other teleosts. Four species of cichlids and three species of wrasses (« Cichlids+Wrasses »: *Choerodon anchorago*, *Labroides dimidiatus*, *Thalassoma hardwicke, Amatitlania nigrofasciata, Maylandia zebra, Neolamprologus brichardi, Ophtalmotilapia boops*, n=26) were compared with three species of teleosts of various orders (« Outgroup »: *Astyanax* mexicanus, *Danio rerio, Salmo trutta*, n=14). (A-E) Comparison of the relative mass of the Tel (A), TeO (B), rFM (C), Cb (D), and rH (E). Cichlids+Wrasses have a relatively larger Tel and rFM compared to the outgroup. (F-J) Comparison of the relative number of cells in the Tel (F), TeO (G), rFM (H), Cb (I), and rH (J). Cichlids+wrasses don’t have a larger proportion of cells in the Tel, but a larger proportion in the rFM compared to the outgroup. Statistical analysis: Mann-Whitney’s test. Each point represents individual values. ns: non significant, *p<0.05, ****p<0.0001. Brain regions: Cb: cerebellum, rFM: rest of the forebrain/midbrain; rH: rest of the hindrain; Tel: telencephalon; TeO: optic tectum.

**Figure 5-figure supplement 3.**
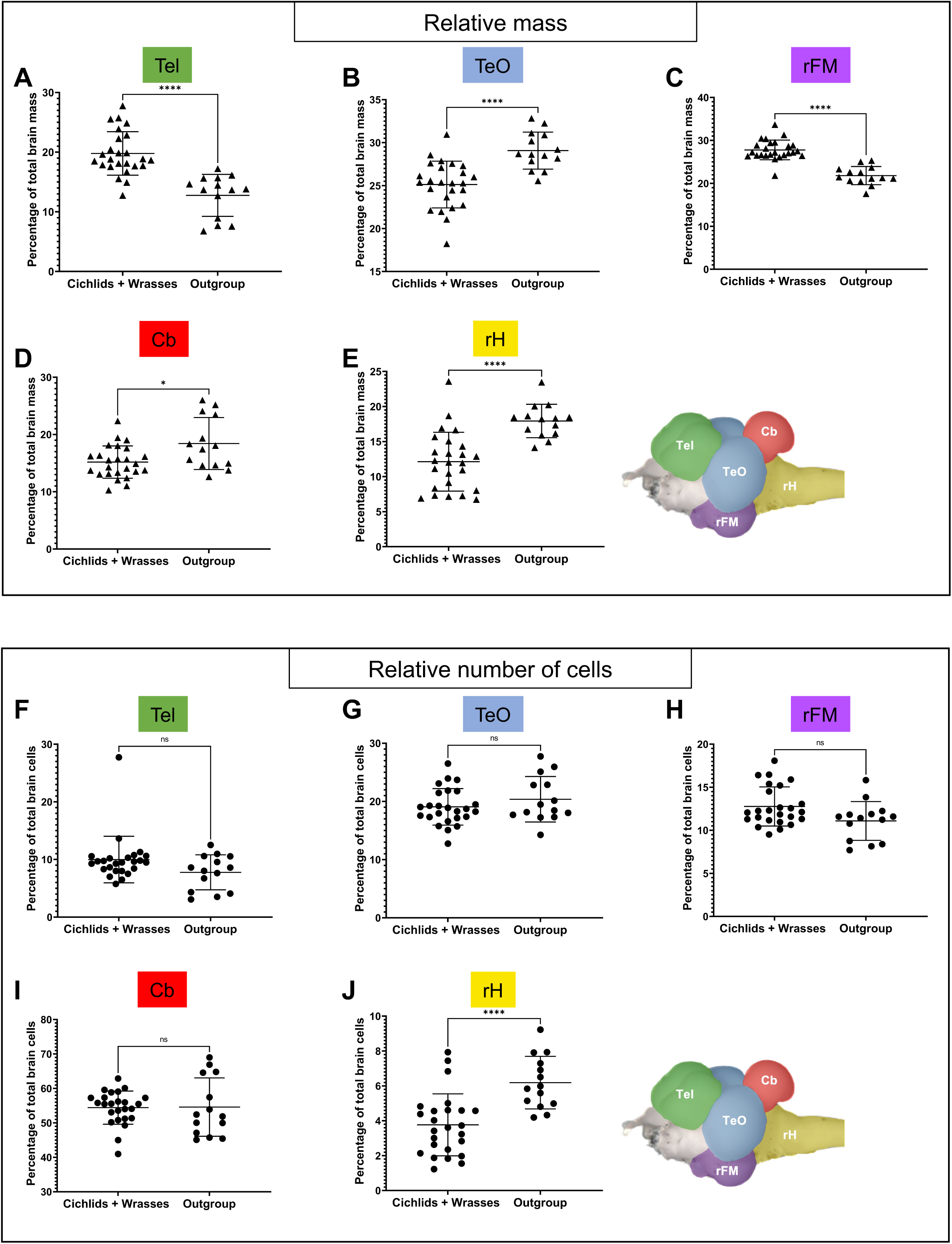
Wrasses and cichlids have a relatively larger telencephalon (Tel) and rest of the forebrain/midbrain (rFM) without a corresponding increase in the relative number of cells compared to other teleosts even after removal of the large *Choerodon anchorago* individual from the analysis. Four species of cichlids and three species of wrasses (« Cichlids+Wrasses »: *Choerodon anchorago*, *Labroides dimidiatus*, *Thalassoma hardwicke, Amatitlania nigrofasciata, Maylandia zebra, Neolamprologus brichardi, Ophtalmotilapia boops*, n=25) were compared with three species of teleosts of various orders (« Outgroup »: *Astyanax mexicanus*, *Danio rerio, Salmo trutta*, n=14). (A-E) Comparison of the relative mass of the Tel (A), TeO (B), rFM (C), Cb (D), and rH (E). Cichlids+Wrasses have a relatively larger Tel and rFM compared to the outgroup. (F-J) Comparison of the relative number of cells in the Tel (F), TeO (G), rFM (H), Cb (I), and rH (J). Cichlids+wrasses don’t have a larger proportion of cells in the Tel nor in the rFM compared to the outgroup. Statistical analysis: Mann-Whitney’s test. Each point represents individual values. ns: non significant, *p<0.05, ****p<0.0001. Brain regions: Cb: cerebellum, rFM: rest of the forebrain/midbrain; rH: rest of the hindrain; Tel: telencephalon; TeO: optic tectum. See also Supplementary file 1.

**Figure 6-figure supplement 1.**
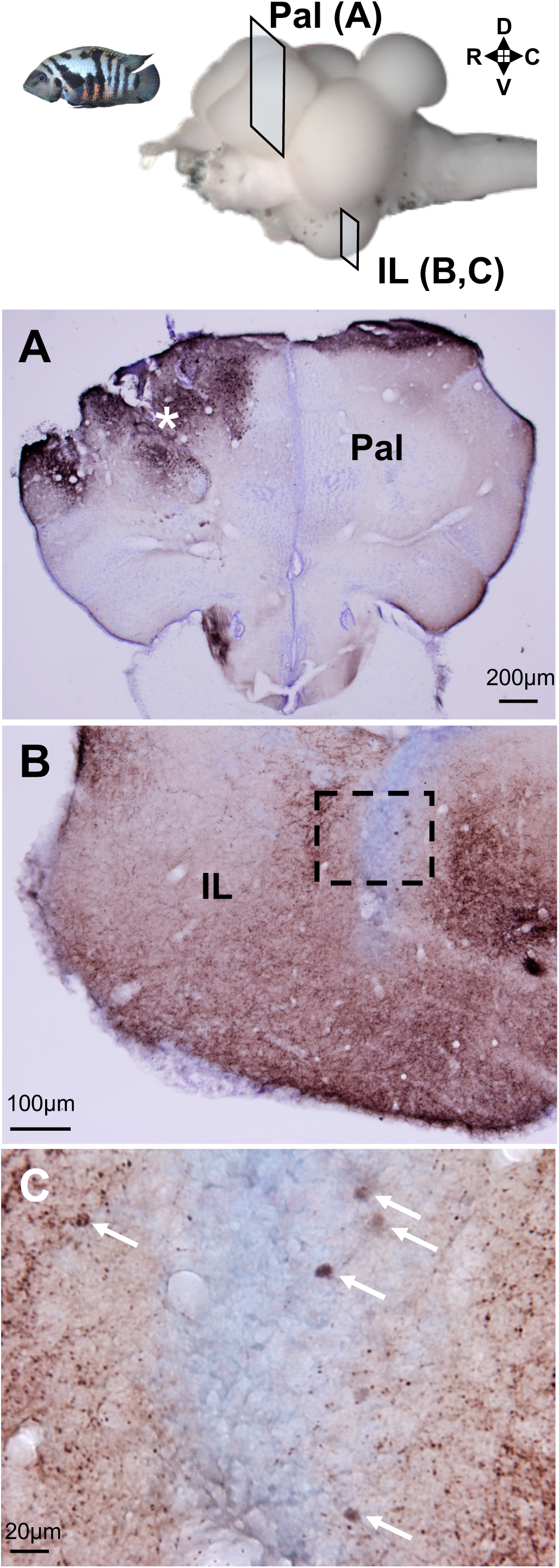
Connectivity of the pallium with the inferior lobe in the cichlid *A. nigrofasciata* visualized using biocytin. Frontal sections showing biocytin tract-tracing in the cichlid *A. nigrofasciata.* The level of the sections is indicated at the top. Brown indicates biocytin, and purple indicates cresyl violet counterstaining. (A) Level of the telencephalon, showing the injection site in the Pal (white asterisk). (B) Ipsilateral IL showing abundant anterogradely labelled fiber terminals (dark brown). The dotted square delineates the area shown in (C). (C) Higher magnification showing the retrogradely labelled cell bodies in IL (white arrows). Brain regions: Pal: pallium, IL: inferior lobe R: rostral; C: caudal; D: dorsal; V: ventral.

