## Supplementary file 1 for "Different ways of evolving tool-using brains in teleosts and amniotes"

### Supplementary file 1: Statistical analysis excluding the large individual of *Choerodon anchorago*

Wrasses in this study were wild caught and tended to be young adults, but one large adult of *Choerodon anchorago* weighing around ten times as much as the other individuals was also sampled. This individual weighed 297.54 g and had a brain mass of 648.9 mg. As this individual diverged from the other 3 *C. anchorago* individuals sampled, we verified whether excluding it from our statistical analysis changed any of our conclusions. The results of this analysis are detailed below.

Teleosts, Primates and Psittacopasserae ($p<0.0001$ in all cases), with the exception of the intercepts of Glires and Primates ($p=0.08$).

Overall, the statistical significance of the results was unchanged when the large *C. anchorago* individual was excluded from the dataset.

Degree of encephalization of sampled species

In order to determine the degree of encephalization of the teleost species sampled in this study, a phylogenetically corrected brain-body allometric slope was estimated using phylogenetically generalized least squares regression test (PGLS) at the Class level on species means of $\log^{10}$ brain and $\log^{10}$ body mass data of the species sampled in this study along with previously published actynopterygians data by Tsuboi et al.( 2018) using RStudio with the CAPER package (Figure 3-figure supplement 1). Residual variance was modelled according to Brownian motion (Felsenstein, 1985) and phylogenetic signal was estimated using Pagel's λ (Lynch, 1991). Phylogenetic relationships between teleost species were based on previously published phylogenetic trees (Rabosky et al., 2013). The phylogenetic regression slope for actinopterygians was of 0.50 ± 0.01 (Adjusted $R^2$: 0.8379, $t=65.891$, $p<0.0001$).

Encephalization was then determined by extracting the residuals of $\log^{10}$-$\log^{10}$ brain and body mass for each species of the dataset to remove allometry in brain size (Sol et al., 2016). The 11 species studied were ranked based on the value of their residual (Table 2). The exclusion of the large *C. anchorago* individual gave a residual of 0.488 for *C. anchorago*, placing it above the wrasse *Thalassoma hardwicke*.

Encephalization and relative mass and relative number of cells of major brain structures

To determine whether there exists a correlation between the degree of encephalization and relative mass and relative number of cells (expressed as the percentage of total brain mass and percentage of total brain cells, respectively) of the five major brain structures dissected, a nonparametric Spearman rank correlation test was used, as there was no way to ascertain the normal distribution of these data. We arranged species by decreasing order of encephalization (Figure 3-figure supplement 3). The test was performed in GraphPad Prism on species means. A significant negative correlation was found between encephalization and the relative mass and relative number of cells in the rH (Figure 3-figure supplement 3E, Spearman r: -0.6970, p=0.0306 and Spearman r: -0.6606, p=0.0438, respectively). No significant correlation with encephalization was found in the four other brain structures for either relative mass or relative number of cells (Figure 3-figure supplement 3A-D, Tel relative mass: Spearman r: 0.5152, p=0.1334; relative number of cells: Spearman r: -0.01818, p=0.973; TeO relative mass: Spearman r: -0.2242, p=0.5367; relative number of cells: Spearman r: 0.09091, p=0.8113; rFM relative mass: Spearman r: 0.3818, p=0.2788; relative number of cells: Spearman r: -0.2242, p=0.5367; Cb relative mass: Spearman r: -0.1758, p=0.6321; relative number of cells: Spearman r: 0.3212, p=0.3679).

Overall, the statistical significance of the results was unchanged when the large *C. anchorago* individual was excluded from the dataset.

<u>Species to species comparison of relative mass, absolute and relative number of cells in major brain structures</u>

Normality of the data was tested using Shapiro-Wilk's test. As normality was not verified for all the species studied, and considering the small sample size, nonparametric Kruskal-Wallis and Dunn's post hoc tests were used to assess the interspecies differences in relative mass, absolute and relative number of cells in the five dissected brain structures. All tests were performed in GraphPad Prism.

Significant differences were found in the absolute number of cells in all five structures (Kruskal-Wallis, $p<0.001$ in all cases). However, post-hoc pairwise comparisons revealed significant differences that were inconsistent across species and brain structures, the only consistently found difference across all structures being between *D. rerio* and *C. anchorago* (Dunn's test, $p<0.05$ in all cases).

Significant differences were found in the relative number of cells in all five structures (Kruskal-Wallis, $p<0.05$ in all cases). However, post-hoc pairwise comparisons revealed significant differences that were inconsistent across species and brain structures.

Significant differences were found in the relative mass of the Tel, TeO, rFM and rH (Kruskal-Wallis, $p<0.01$ in all cases). No significant difference was found between species in the relative mass of the Cb (Kruskal-Wallis, $p=0.0649$). However, post-hoc pairwise comparisons didn't reveal significant differences between species across the four structures, except for a modest difference in the relative mass of the rH between *A. mexicanus* and *T. hardwicke* (Dunn's test, $p=0.0292$).

Overall, the statistical significance of the results was unchanged when the large *C. anchorago* individual was excluded from the dataset.

<u>Comparison of relative mass and relative number of cells in major brain structures based on behavioral repertoire</u>

Among the teleost species studied, wrasses display the most complex behavioral phenotypes. To determine whether this behavioral repertoire is associated with differences in relative mass and relative number of cells in major brain structures compared to other teleosts, the three species of wrasse (*C. anchorago* (n=3), *T. hardwicke* (n=3) and *L. dimidiatus* (n=3)) were grouped together (n=9) and compared to all the other species (*M. zebra* (n=3), *N. brichardi* (n=5), *O. boops* (n=3), *A. nigrofasciata* (n=5), *A. mexicanus* (n=5), *D. rerio* (n=5) and *S. trutta* (n=4), grouped as "Other fish" (n=30)). As normality could not be satisfied for all structures in both groups, a nonparametric Mann-Whitney test was used. Regarding the relative mass, wrasses had a significantly larger Tel and rFM compared to other teleosts (Mann-Whitney's test, $p<0.0001$ and $p=0.008$, respectively), and a significantly smaller Cb and rH ($p=0.0035$ and $p<0.0001$, respectively). No significant differences were found in the relative mass of the TeO ($p=0.8571$). Regarding the relative number of cells, wrasses had a significantly lower relative number of cells in the rH compared to the other teleosts (Mann-Whitney's test, $p<0.0001$). No significant differences were found in the relative number of cells of the other four structures (Tel: $p=0.1226$; TeO: $p=0.1494$; rFM: $p=0.0636$; Cb: $p=0.9088$). Normality and Mann-Whitney tests were performed in GraphPad Prism.

Overall, the statistical significance of the results was unchanged when the large *C. anchorago* individual was excluded from the dataset.

<u>Comparison of relative mass and relative number of cells in major brain structures based on phylogeny</u>

In order to assess the differences in relative mass and relative number of cells in the five dissected brain structures based on phylogeny, species were grouped into different clusters: *M. zebra* (n=3), *N. brichardi* (n=5), *O. boops* (n=3) and *A. nigrofasciata* (n=5), all members of the Cichlidae family, were grouped as "cichlids" (n=16). *C. anchorago* (n=3), *T. hardwicke* (n=3) and *L. dimidiatus* (n=3), members of the Labridae family, were grouped as “wrasses” (n=9). *A. mexicanus* (n=5) (Characidae), *D. rerio* (n=5) (Cyprinidae) and *S. trutta* (n=4) (Salmonidae) being phylogenetically distant, were grouped together as an “outgroup” (n=14) because of the n=1 species per family sample size.

Cichlids and wrasses are part of the suborder Labrodei and are very closely phylogenetically related. Cichlids and wrasses were grouped together as “Cichlids+Wrasses” (n=25) and compared to the outgroup using a nonparametric Mann-Whitney test, as normality could not be satisfied for all structures in both groups (Figure 5-figure supplement 3). Regarding the relative mass, Cichlids+Wrasses had a significantly larger Tel (Figure 5-figure supplement 3A) and rFM (Figure 5-figure supplement 3C) compared to the outgroup (Mann-Whitney’s test, $p<0.0001$ in both structures), and a significantly smaller TeO (Figure 5-figure supplement 3B), Cb (Figure 5-figure supplement 3D) and rH (Figure 5-figure supplement 3E) ($p<0.0001$, $p=0.0301$ and $p<0.0001$, respectively). Regarding the relative number of cells, Cichlids+Wrasses had a significantly lower relative number of cells in the rH (Figure 5-figure supplement 3J) compared to the Outgroup (Mann-Whitney’s test, $p<0.0001$). No significant differences were found in the relative number of cells of the other four structures (Figure 5-figure supplement 3F-I, Tel: $p=0.1489$; TeO: $p=0.4088$; rFM: $p=0.0625$; Cb: $p=0.6127$). All tests were performed in GraphPad Prism.

Excluding the large individual of *C. anchorago* from the dataset led to the relative number of cells in the rFM for Cichlids+Wrasses to become non-significant compared to the Outgroup.

#### Supplementary Information references

Felsenstein, J. (1985). Phylogenies and the Comparative Method. *The American*

*Naturalist*, *125*(1), 1-15.

Herculano-Houzel, S., Catania, K., Manger, P. R., & Kaas, J. H. (2015). Mammalian Brains Are Made of These : A Dataset of the Numbers and Densities of Neuronal and Nonneuronal Cells in the Brain of Glires, Primates, Scandentia, Eulipotyphlans, Afrotherians and Artiodactyls, and Their Relationship with Body Mass. *Brain, Behavior and Evolution*, *86*(3-4), 145-163. https://doi.org/10.1159/000437413

Lynch, M. (1991). METHODS FOR THE ANALYSIS OF COMPARATIVE DATA IN EVOLUTIONARY BIOLOGY. *Evolution; International Journal of Organic Evolution*, *45*(5), 1065-1080. https://doi.org/10.1111/j.1558-5646.1991.tb04375.x

Olkowicz, S., Kocourek, M., Lučan, R. K., Porteš, M., Fitch, W. T., Herculano-Houzel, S., & Němec, P. (2016). Birds have primate-like numbers of neurons in the forebrain. *Proceedings of the National Academy of Sciences*, *113*(26), 7255-7260. https://doi.org/10.1080/19419899.2013.835743

Rabosky, D. L., Santini, F., Eastman, J., Smith, S. A., Sidlauskas, B., Chang, J., & Alfaro, M. E. (2013). Rates of speciation and morphological evolution are correlated across the largest vertebrate radiation. *Nature Communications*, *4*, 1958. https://doi.org/10.1038/ncomms2958

Sol, D., Sayol, F., Ducatez, S., & Lefebvre, L. (2016). The life-history basis of behavioural innovations. *Philosophical Transactions of the Royal Society of London. Series B, Biological Sciences*, *371*(1690), 20150187. https://doi.org/10.1098/rstb.2015.0187

Tsuboi, M., van der Bijl, W., Kopperud, B. T., Erritzøe, J., Voje, K. L., Kotrschal, A., Yopak, K. E., Collin, S. P., Iwaniuk, A. N., & Kolm, N. (2018). Breakdown of brain–body allometry and the encephalization of birds and mammals. *Nature Ecology and Evolution*, *2*(9), 1492-1500. https://doi.org/10.1038/s41559-018-0632-1

Warton, D. I., Duursma, R. A., Falster, D. S., & Taskinen, S. (2012). Smatr 3– an R package for estimation and inference about allometric lines. *Methods in Ecology and Evolution*, *3*(2), 257-259. https://doi.org/10.1111/j.2041-210X.2011.00153.x
